# Macromolecular crowding and supersaturation protect hemodialysis patients from the onset of dialysis-related amyloidosis

**DOI:** 10.1101/2022.02.01.478730

**Authors:** Kichitaro Nakajima, Keiichi Yamaguchi, Masahiro Noji, César Aguirre, Kensuke Ikenaka, Hideki Mochizuki, Lianjie Zhou, Hirotsugu Ogi, Toru Ito, Ichiei Narita, Fumitake Gejyo, Hironobu Naiki, Suguru Yamamoto, Yuji Goto

## Abstract

Dialysis-related amyloidosis (DRA), a serious complication among long-term hemodialysis patients, is caused by amyloid fibrils of β_2_-microglobulin (β2m). Although high serum β2m levels and a long dialysis vintage are the primary and secondary risk factors for the onset of DRA, respectively, patients with these do not always develop DRA, indicating that there are additional risk factors. To clarify these unknown factors, we investigated the effects of human sera on β2m amyloid fibril formation. Although sera markedly inhibited amyloid fibril formation, the inhibitory effects were weaker for sera collected from dialysis patients than for control sera. When sera collected before and after maintenance dialysis treatments were compared, the latter inhibited amyloid fibril formation more than the former. These results indicate that, although the inhibitory effects of sera were deteriorated in long-term dialysis patients, they were ameliorated by maintenance dialysis treatments in the short term. Maintenance dialysis decreases the body weight by 5% and consequently increases the concentrations of serum components. Among these components, we found that serum albumin prevented amyloid fibril formation based on macromolecular crowding effects, and that the decreased serum albumin concentration in dialysis patients is a tertiary risk factor for the onset of DRA. The model was constructed assuming accumulative effects of three risk factors and may be useful for predicting the onset of DRA. Furthermore, the model suggested the importance of monitoring temporary and accumulated risks to prevent the development of amyloidoses in general, which occurs based on supersaturation-limited amyloid fibril formation in a crowded milieu.

## Introduction

Dialysis-related amyloidosis (DRA), one of the systemic amyloidoses ^1,2^, is a serious complication of long-term hemodialysis therapy and a threat to the health of dialysis patients ^3–8^. The pathological hallmark of DRA is the deposition of amyloid fibrils of β_2_-microglobulin (β2m) in the peritenons and synovial membranes of the carpal tunnel. β2m is a component of the major histocompatibility complex (MHC) class I expressed on the surface of all nucleated cells ^9^. β2m monomers are released into bloodstream when MHCs dissociate. In individuals with healthy kidneys, most of the released β2m monomers are eliminated from the bloodstream by glomerular filtration and subsequent reabsorption and catabolism by the proximal tubules, keeping the serum β2m concentration at ~1 μg/mL. However, in long-term hemodialysis patients, the concentration of β2m increases up to ~50 μg/mL because it is not degraded due to kidney failure and is not completely eliminated by maintenance dialysis treatments. Consequently, β2m forms amyloid fibrils and causes DRA in patients who have been on hemodialysis therapy for more than 10 years. These facts indicate that a high serum β2m concentration and long dialysis vintage are the primary and secondary risk factors, respectively, for the onset of DRA. However, patients with these risks do not always develop DRA ^4–8^, indicating that there are additional risk factors (Figure S1).

In Japan, dialysis technology has progressed markedly in the past 40 years and has extended the time for first-time carpal tunnel surgery (CTS), as a proxy for the onset of DRA ^6,8^. A comparison of cohorts of chronic hemodialysis patients (more than 200,000 in total) in 1998 and 2010 showed that the risk for DRA decreased from 1998 to 2010 ^6^. The risk of first-time CTS was almost halved, even though it was still 10% in the 2010 cohort with a dialysis vintage longer than 20 years. The survey indicated that improvement in dialysis technology, particularly, the introduction of biocompatible dialysis membranes, contributed to the decreased risk ^6^. Recently, a retrospective survey of 222 patients across 4 periods from 1982 to 2019 showed that improvement of β2m clearance via advances in dialysis technology contributed to extending the time between starting hemodialysis and the first surgery for CTS ^8^. The time was 12.4 years in period 1 (1982-1989) but extended to 22.4 years in period 4 (2010-2019).

Among various amyloidogenic proteins, β2m, a β-barrel globular protein consisting of 99 amino acid residues, is particularly useful for addressing the relationship between protein folding and misfolding (i.e., amyloid fibril formation). We have studied β2m amyloid fibril formation ^10–15^ from the physicochemical viewpoint that amyloid fibril formation is similar to crystallization of solutes ^13,16–18^. According to this view, amyloid fibrils are crystal-like aggregates of denatured proteins, which are formed above solubility upon breaking supersaturation. Supersaturation, a fundamental phenomenon of nature with underlying physicochemical mechanisms remaining elusive ^19–21^, is required for crystallization and is involved in numerous phenomena, e.g., the supercooling of water prior to ice formation or various types of lithiasis caused by small organic compounds ^13^. The same will be true for crystal-like amyloid fibrils. Under supersaturated conditions above solubility, an unknown trigger can break supersaturation, leading to amyloid fibril formation ^11,13,22^.

To gain insights into the pathogenesis of DRA under macromolecular crowding, we studied the effects of serum components on β2m amyloid fibril formation. It should be noted that, although previous studies ^23–26^ investigated amyloid fibril formation in a crowded milieux (e.g., serum milieu), there is no consensus on the effects of macromolecular crowding in a serum milieux. With HANABI-2000 ^27–29^, an ultrasonication-forced amyloid fibril inducer, the effects of sera from hemodialysis patients on β2m amyloid fibril formation were investigated. To perform a systematic investigation, we collected over 100 sera from individuals with or without hemodialysis therapy. The results showed that sera inhibited β2m amyloid fibril formation based on the macromolecular crowding effects arising from serum albumin, the most abundant protein in sera, and that the inhibitory effects were weaker for hemodialysis patients, particularly before maintenance dialysis treatments, because of the decreased serum albumin concentration. We modelled the development of DRA assuming that the primary, secondary, and tertiary risk factors are an increased β2m concentration, a long dialysis vintage, and decreased serum albumin concentration, respectively. The model suggested that accumulated risks break the extracellular proteostasis network ^30^ under supersaturation, leading to amyloid fibril formation and thus the development of amyloidosis. The current model based on solubility- and supersaturation-limited amyloid fibril formation will be applicable to a variety of amyloidoses and, furthermore, will be useful for devising therapeutic strategies.

## Results

### HANABI assay for β2m amyloid fibril formation

Previously, Noji *et al*. ^14,15^ reported that β2m monomers even at a neutral pH form amyloid fibrils at high temperature under agitation, effective conditions to break supersaturation. Thus, control reactions were performed at 1.0 mg/mL β2m at pH 7.4 and 60 °C. To apply further effective agitation to induce amyloid fibril formation, we adopted an originally developed ultrasonic instrument, HANABI-2000 ^27–29^, which accelerates amyloid fibril formation ^10,11,31,32^ by the effects of ultrasonic cavitation ^33,34^ and monitors amyloid formation by amyloid-specific fluorescence dye, thioflavin-T (ThT) ^35^. Scheme of the experiment is described in Figure S2. Consistent with previous reports ^14,15^, β2m solutions did not show an increase in ThT fluorescence intensity without ultrasonication, but did when ultrasound was applied (Figure 1A), indicating amyloid formation under ultrasonication.

**Figure 1.**
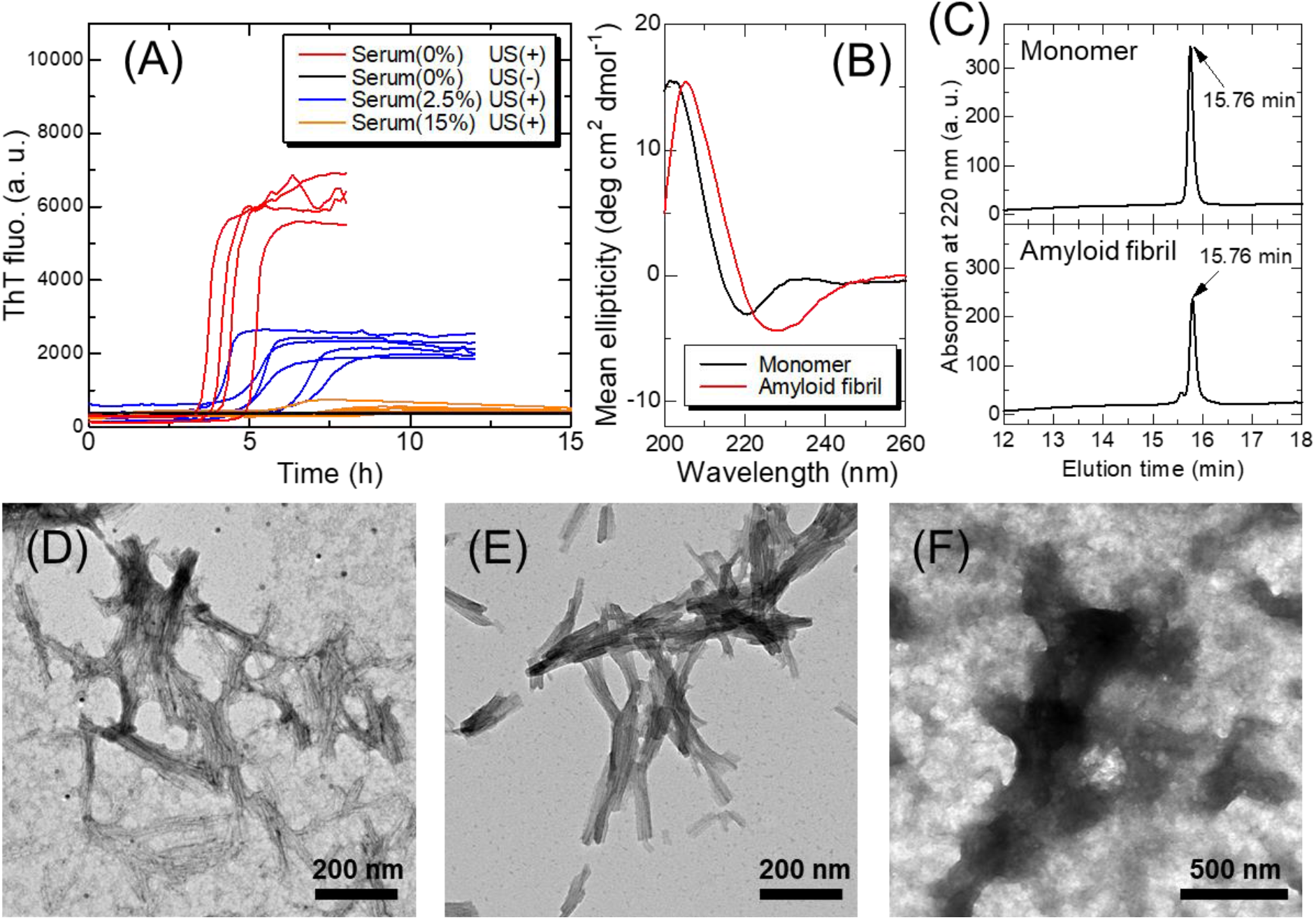
(A) ThT fluorescence kinetics of β2m solutions at various conditions (*n* ≥ 4). Serum was collected from a non-dialysis control #9 (DT(-)). (B) CD spectra of the initial monomers and amyloid fibrils formed by ultrasonication. (C) High performance reversed-phase chromatograms for intact monomers (upper) and amyloid fibrils formed by ultrasonication (lower). TEM images of (D) ThT-positive aggregates formed by ultrasonication, (E) aggregates seeded and elongated from the amyloid fibrils formed by ultrasonication, and (F) ThT-negative aggregates formed in the presence of 15% (v/v) serum.

After the ultrasonic experiments, the aggregates formed were analyzed. The circular dichroism (CD) spectra showed a transition from native monomers with a minimum at 220 nm to amyloid fibrils with a minimum at 230 nm (Figure 1B), consistent with previous reports ^12,14,15^. High performance reversed-phase chromatography revealed the persistence of intact monomers (Figure 1C, SI Result 1, and Figure S3), confirming that no fragmentation or other damage of β2m monomers occurred during the ultrasonic experiments. Transmission electron microscopy (TEM) images of aggregates also indicated a fibrous morphology (Figure 1D, SI Result 2, and Figure S4). However, since TEM images of amyloid fibrils produced under ultrasonication often showed poor contrast because of the extensive fragmentation ^11,36^ and unknown effects, we performed a seeding reaction to prepare longer amyloid fibrils. The seeded amyloid fibrils clearly showed a fibrous morphology with improved contrast (Figure 1E, SI Result 2, and Figure S4).

To investigate the effects of human sera on amyloid fibril formation, we then added an aliquot of sera collected from a non-dialysis control without dialysis therapy (DT(-), #9) to the standard β2m solution (Figure 1A). At a serum concentration of 2.5% (v/v), although a ThT fluorescence increased, the lag time was much longer, and the final ThT fluorescence intensity was low. At a serum concentration of 15% (v/v), ThT fluorescence remained unchanged within 15 h. TEM images without a clear fibrous morphology suggested the formation of amorphous aggregates (Figure 1F). These results indicate that the addition of sera inhibited β2m amyloid fibril formation.

### Concentration-dependent inhibitory effects of sera

To systematically investigate the inhibitory effects of sera on β2m amyloid fibril formation, three types of sera were prepared: control sera from individuals without dialysis therapy (DT(-),#9), and sera collected from dialysis patients immediately before (DT(+, Pre), #8) and after (DT(+, Post), #8) maintenance dialysis treatments. All dialysis patients examined in this study underwent maintenance dialysis treatments involving 4-5 h sessions conducted three times a week (SI Methods). The effects of sera on amyloid formation were assayed at varying serum concentrations, 0.3-15% (v/v) (Figure 2A-C). Here, β2m monomers from sera were less than 1% of the recombinant β2m monomers in the standard solution (1.0 mg/mL), indicating that the effects of β2m monomers present in sera were negligible (SI Result 3 and Figure S5). In other words, we eliminated the primary risk factor to focus on other risk factors. The kinetics monitored by ThT fluorescence were analyzed using two parameters, a lag time and the maximum intensity of ThT fluorescence, where the lag time was when the ThT fluorescence intensity became one tenth of the maximum.

**Figure 2.**
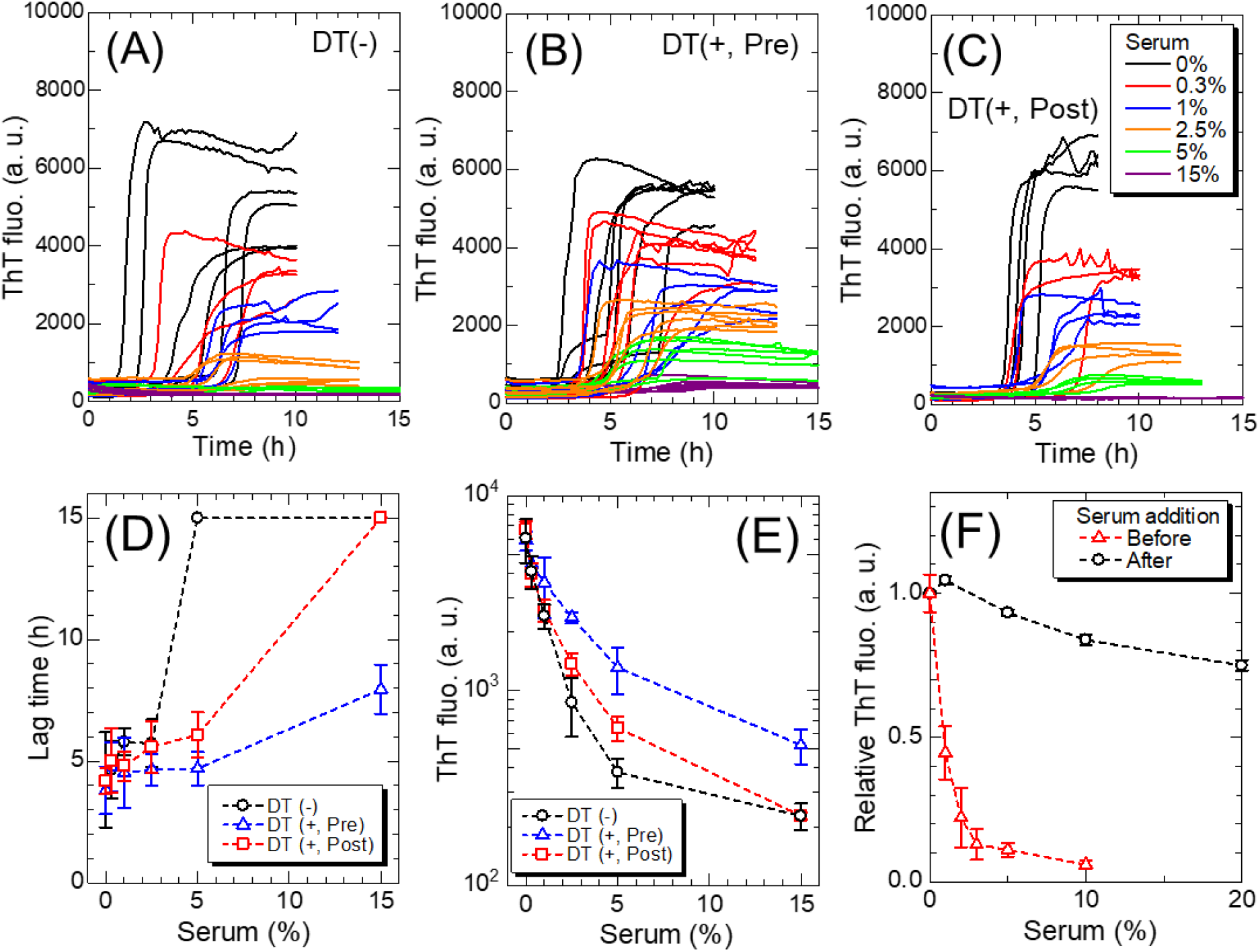
ThT fluorescence kinetics at varying concentrations of sera collected from (A) a non-dialysis control (DT(-), #9) and a dialysis patient immediately (B) before (DT(+, Pre), #8) and (C) after maintenance dialysis treatment (DT(+, Post), #8), respectively (*n* ≥ 4). Dependence of (D) the lag time and (E) ThT fluorescence intensity on serum concentration for each serum. (F) Decrease in the ThT fluorescence intensity with the addition of serum to the sample solution before and after amyloid fibril formation. Error bars denote the standard deviation.

For all types of sera, the higher the concentration of sera added, the lower the lag time (Figure 2D) and longer the ThT fluorescence intensity (Figure 2E). It was initially unclear whether the lower ThT intensity represents a decreased amount of amyloid fibrils or an apparent decrease due to optical absorption of incident beams by serum components, etc. To investigate the relationship between ThT fluorescence intensity and the amount of amyloid fibrils, a series of seeding experiments were conducted using products obtained at various serum concentrations (SI Result 4 and Figure S6). The lag time of the ThT kinetics verified by the seeding experiments showed an inverse correlation with the initial ThT fluorescence intensities of the seed solution. Moreover, when serum was added to the preformed amyloid fibrils, the decrease in the ThT fluorescence intensity due to optical absorption by serum components was much less than the decrease caused by amyloid fibril formation in the presence of serum (Figure 2F). These results indicate that, even in the presence of serum, the maximal ThT fluorescence intensity was proportional to the amount of amyloid fibrils, and that the decrease in ThT fluorescence maximum intensity represented the inhibitory effects of serum components.

### Deterioration of inhibitory effects by long-term dialysis

To focus on the effects of dialysis vintage, we examined the difference in the effects of sera with and without hemodialysis therapy for a total of 60 sera at different dialysis vintages (Figure S7). As shown in Figure 3A, 30 of the 60 sera were collected from dialysis patients (DT(+, Pre), *N* = 30), and the remaining 30 sera were collected from non-dialysis controls (DT(-), *N* = 30). The sera from dialysis patients were collected before maintenance dialysis treatments, and the serum concentration added was 5% (v/v), where the serum-dependent differences in the lag time and ThT fluorescence intensity were large (Figure 2D,E). The representative kinetics are shown in Figure 3B. Without serum addition, the ThT kinetics indicated rapid amyloid fibril formation with a lag time of approximately 2 h and a marked increase in the ThT fluorescence intensity, confirming the rapid formation of a large amount of amyloid fibrils. Although the two types of sera markedly inhibited amyloid formation regarding both the lag time and ThT fluorescence intensity, the degree of inhibition was different between the two groups. The average lag times were 3.9 and 5.4 h for DT(+, Pre) and DT(-) groups, respectively (Figure 3C), and the maximal ThT fluorescence intensities were 6600 and 2300 for DT(+, Pre) and DT(-) groups, respectively (Figure 3D). These results suggest that, even after eliminating the primary risk of an increased β2m concentration, sera from dialysis patients retained an additional risk of amyloid fibril formation in terms of reducing serum-dependent inhibitory effects.

**Figure 3.**
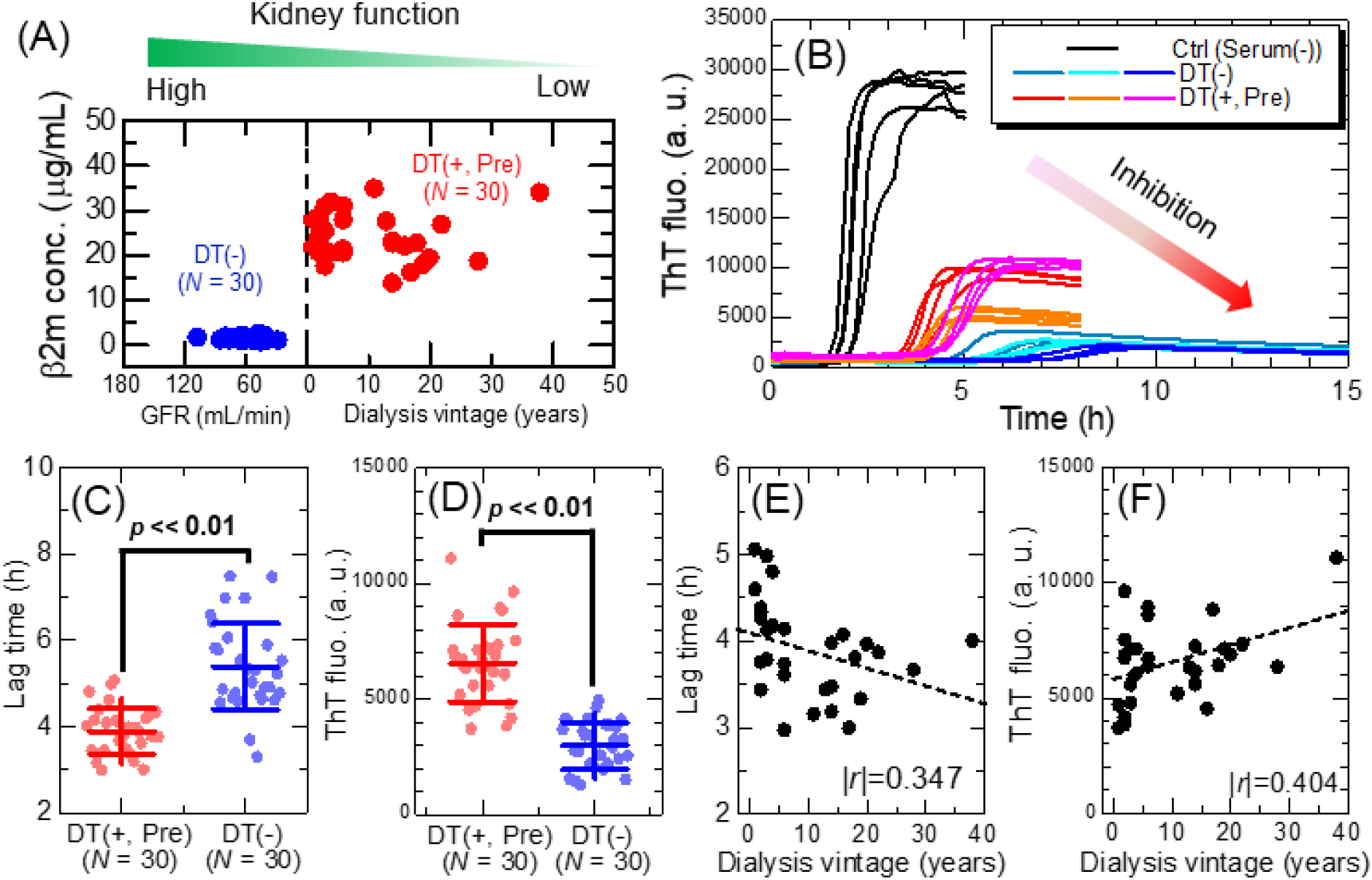
(A) Clinical characteristics of non-dialysis control (DT(-), *N* = 30) and dialysis patient (DT(+, Pre), *N* = 30) groups. For the non-dialysis group, their glomerular filtration rates (GFR) are used as an index of kidney function, because they have never received dialysis treatments. (B) Representative ThT fluorescence kinetics using sera collected from non-dialysis controls and dialysis patients. For each sample, measurement was performed multiple times for independent solutions (*n* ≥ 4). Comparison of the effect of sera with or without dialysis treatment on β2m fibril formation in terms of the (C) lag time and (D) ThT fluorescence intensity. For all 60 samples, row data are shown in Figure S7. Dialysis-vintage dependence of (E) the lag time and (F) ThT fluorescence intensity (*N* = 30).

A previous study ^37^ showed that the secondary risk factor for the onset of DRA, following the primary risk factor of an increased serum β2m level, is a long dialysis vintage ^4–6,8^. The relationship between the dialysis vintage and lag time (Figure 3E) or ThT fluorescence intensity (Figure 3F) revealed a minimal correlation between them. The results suggest that, although the temporal risks, other than the increased β2m concentration, notably increased in dialysis patients, they did not change markedly during the dialysis vintage. Given that the dialysis vintage is a strong risk factor for the onset of DRA, accumulated (or integrated) temporal risks over the dialysis vintage may determine the onset of DRA. In other words, the results of HANABI assays provide an indicator of the temporal risks of DRA onset. It is likely that a combination of temporal risks, an increased β2m concentration, and dialysis vintage collectively provides a susceptibility risk biomarker ^38^ for DRA, which represents the potential for the onset of disease in an individual who does not currently have clinically apparent symptoms.

### Amelioration of inhibitory effects by maintenance dialysis

Next, we examined the effects of maintenance dialysis treatments, conducted three times a week, on the inhibitory effects of serum. As shown in Figure 4A, maintenance dialysis treatment markedly changes the serum milieu: the β2m monomers are eliminated approximately more than 50%, and body weights of patients are decreased because of the expulsion of water. Sera were collected from a total of 28 dialysis patients immediately before (DT(+, Pre)) and after (DT(+, Post)) maintenance dialysis treatments, and HANABI assays were conducted. The patients were different from those used to examine the effects of dialysis vintage. Notably, the maintenance dialysis treatments markedly recovered the inhibitory effects of serum against amyloid formation (Figure 4B). Statistical analysis (SI Result 5 and Figure S8) confirmed the ameliorating effects in terms of a longer lag time (Figure 4C) and lower ThT fluorescence intensity (Figure 4D). These results indicate that, although the inhibitory effects of sera were deteriorated in long-term dialysis patients, they were ameliorated by maintenance dialysis treatments in a short term.

**Figure 4.**
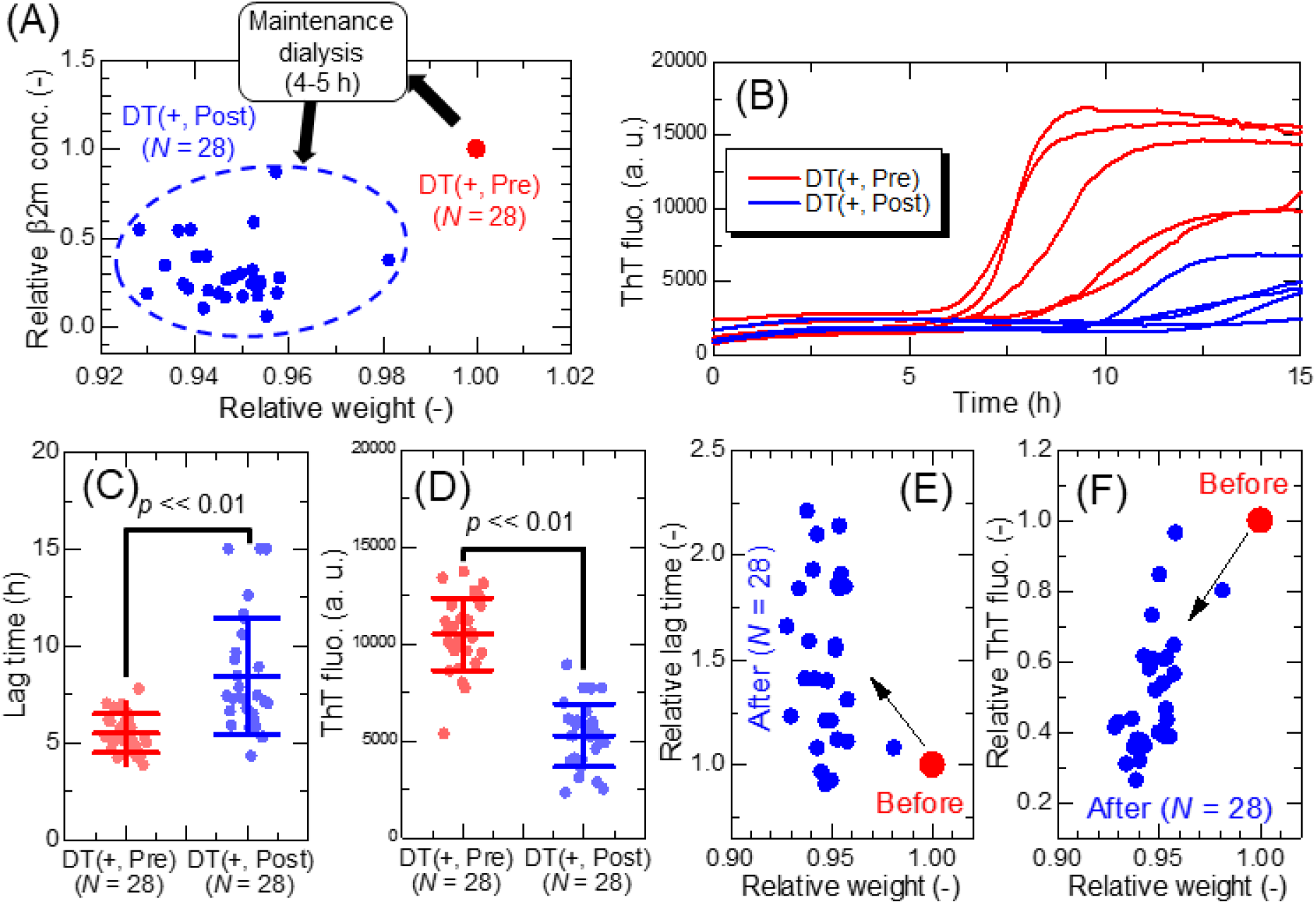
(A) Effects of maintenance dialysis treatment on the serum β2m concentration and body weight of dialysis patients (*N* = 28), who donated their serum immediately before (DT(+, Pre)) and after (DT(+, Post)) maintenance dialysis treatment. The relative changes in the two parameters were shown with respect to their values before dialysis treatment. (B) ThT fluorescence kinetics of β2m fibril formation with 5% (v/v) addition of sera collected from an identical dialysis patient immediately before (DT(+, Pre), #15) and after (DT(+, Post), #15) a single maintenance dialysis treatment. Significance examination for the effects of serum before and after dialysis treatment in terms of the (C) lag time and (D) ThT fluorescence intensity (*N* = 28). For all dialysis patients, row data are shown in Figure S8. Correlation between change in relative weight and (E) lag time and (F) ThT fluorescence intensity.

Importantly, a single maintenance dialysis treatment eliminates a large amount of extra water (usually ~5% of body weight) from a patient’s body, largely from blood, along with biological waste, playing the role of the kidney in patients with end-stage renal failure. The weight change of patients studied in the present study ranged from 2 to 8% (Figure 4E,F). Although plasma refilling occurs to maintain the blood volume during dialysis treatments ^39^, the concentrations of serum components transiently and immediately increase after maintenance dialysis due to the expulsion of water. This is likely to recover the inhibitory effect, resulting in a decrease in the temporary risk (*TR*) (see Discussion).

### Serum albumin is primarily responsible for the inhibitory effects

To reveal the underlying mechanism of the change in the inhibitory effects of sera, correlations between concentrations of 27 serum components and amyloidogenicities in terms of the lag time or ThT fluorescence intensity were systematically investigated (Figure S9). We included data for both calculation of the Pearson correlation coefficient between the concentration of each serum component and the lag time or ThT fluorescence intensity. The correlations suggested three candidates of serum components that markedly affect amyloid fibril formation: serum albumin, ureic acid, and creatinine (SI Result 6). We separately investigated the effects of the three candidates, revealing that only serum albumin significantly inhibited amyloid formation (Figure S10). Lower albumin concentrations tended to induce faster kinetics and larger amounts of amyloid fibrils (Figure S9, panel 23), indicating that serum albumin plays a major role in inhibiting β2m amyloid formation in a serum milieu.

To address amyloid fibril formation in a crowded milieu, we performed HANABI-2000 assays in the presence of 5 mg/mL serum albumin or 5 mg/mL polyethylene glycol (PEG), a control macromolecular crowder (Figure 5A-C). The results showed that, although serum albumin markedly inhibited amyloid formation, PEG slightly extended the lag time, but the resulting amount of amyloid fibrils monitored by ThT fluorescence slightly increased.

**Figure 5.**
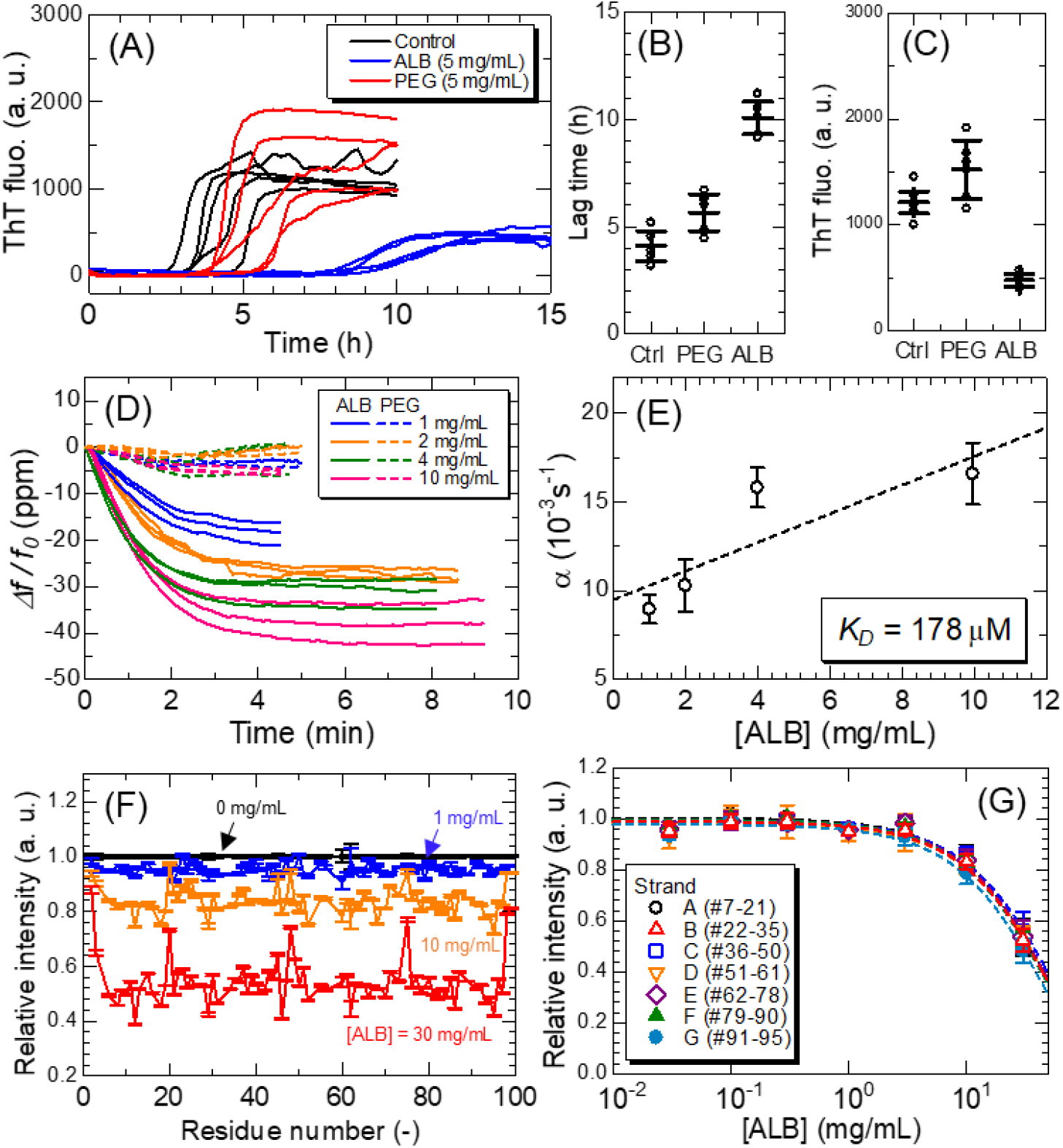
(A-C) The effects of serum albumin or polyethylene glycol on β2m amyloid fibril formation. (A) ThT fluorescence kinetics in the presence of 5 mg/mL serum albumin or polyethylene glycol (*n* = 5), (B) their lag times, and (C) ThT fluorescence intensities. Error bars denote the standard deviation. (D,E) Interactions between β2m monomer and serum albumin monitored by QCM experiments. (D) Relative frequency change (*n* = 3). (E) Calculation of the dissociation constant, *K*_D_, between β2m monomer and serum albumin. (F) Relative intensity changes in the NMR signal of the ^15^N-labeled β2m monomer at different albumin concentrations. (G) Relative intensity changes in the NMR signals averaged over each strand.

Interactions between β2m monomers and the two crowders were investigated by wireless-electrodeless quartz crystal microbalance (QCM) biosensor ^40,41^ and solution nuclear magnetic resonance (NMR) measurements. In QCM analysis, serum albumin bound to β2m monomer in a concentration-dependent manner, but PEG did not (Figure 5D). We considered that the simplest mechanism of complex formation existed between β2m monomers and serum albumin with a stoichiometry of 1:1. From the fitting analysis of frequency-change curves with a function Δ*f*/*f*_0_ = *A*(*e*^−*αt*^ – 1) ^42^, where *A* and *α* are fitting constants, the obtained *α* values were plotted against the albumin concentration (Figure 5E). Because the *α* value is written as α = *k*_a_[ALB] + *k*_d_ ^42^, the *k*_a_ and *k*_d_ values, which are the rate constants for association and dissociation reactions, respectively, can be obtained from the slope and intercept of the linear regression line in Figure 5E, respectively. The dissociation constant, *K*_D_ = *k*_d_/*k*_a_, was calculated as *K*_D_ = 178 μM.

NMR analysis showed the decrease in the NMR signal intensity of ^15^N-labeled β2m monomers with an increasing serum albumin concentration (Figure 5F). Since the β2m native structure consists of seven β-strands, the averaged relative intensity changes were calculated for respective β-strands and compared (Figure 5G), indicating that there is no specific binding site in the β2m monomers. The dissociation constant estimated from the decrease in the NMR signal was ~500 μM, being on the same order as that obtained from QCM analysis. These results indicate that the interaction between β2m monomer and serum albumin is weak binding ^43^.

The effects of molecular crowders on amyloid fibril formation are explained by three dominant mechanisms ^23,24,44^: (i) volume exclusion effects, accelerating amyloid fibril formation ^25^; (ii) interactions with crowders, decelerating amyloid fibril formation ^26^; and (iii) decrease in the diffusion constant, decelerating amyloid fibril formation ^45^. The results showed that serum albumin inhibited β2m amyloid formation by mechanism (ii): Weak interactions between serum albumin and β2m monomer decreased the concentration of unfolded β2m through the “law of mass action” ^14^. On the other hand, PEG molecules were inert to β2m monomer, slightly increasing the amount of amyloid fibrils by mechanism (i), but they retarded the kinetics by mechanism (iii). Serum albumin was primarily responsible for the inhibitory effects despite the weak interactions. This could be achieved by a huge amount of serum albumin in a serum milieu: The concentration of serum albumin was ~45 mg/mL, being the highest among serum proteins with a total concentration of ~70 mg/mL.

If serum albumin possesses the ability to bind to serum proteins with a high affinity, most of them would fail to exert their biological functions because of strong trapping by serum albumin. Thus, it is inferred that the huge amount and weak affinity of serum albumin is crucial for keeping the proteostasis network in a serum milieu, and that a decrease in the serum albumin concentration may be a risk factor for not only DRA, but also other systemic amyloidoses.

## Discussion

### Model of β2m amyloid fibril formation in serum milieu

#### Overall schemes

Focusing on the inhibitory effects of serum albumin, we created a model of β2m amyloid formation in a serum milieu (Figure 6A). We considered that the simplest stoichiometric binding existed between serum albumin (ALB) and β2m monomer in its native state (N) (Scheme 1), a two-state unfolding/folding between N and denatured state (D) (Scheme 2), and the seed-dependent amyloid elongation mechanism, where P represents polymeric amyloid fibrils (Scheme 3).

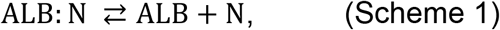

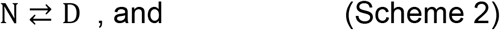

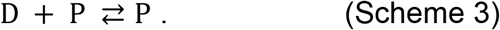

**Figure 6.**
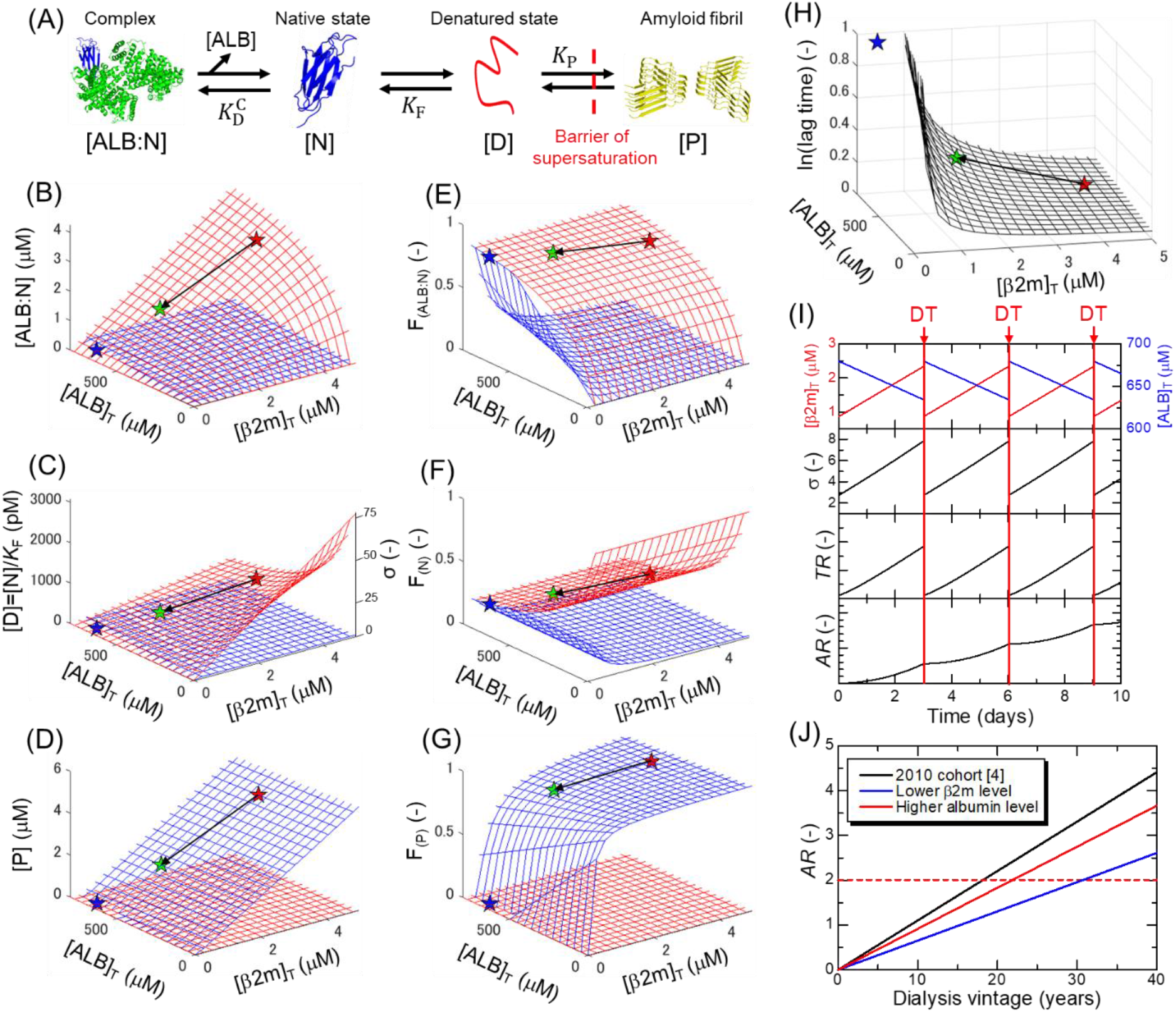
(A) Schematic illustration of amyloid fibril formation in a serum milieu. (B-D) Dependences on total β2m ([β2m]_T_) and albumin ([ALB]_T_) concentrations of (B) native β2m-serum albumin complex, (C) denatured monomer, and (D) amyloid fibrils, before (red mesh) and after (blue mesh) breakdown of supersaturation. In panel c, the axis of the degree of supersaturation, *σ* = [D]^S^/[D]_C_, is also shown. (E-G) Dependences as shown in panels B-D are represented by fractions of (E) native β2m-serum albumin complex, (F) native monomer, and (G) amyloid fibrils. (H) The relative change in the lag time as a function of [β2m]_T_ and [ALB]_T_. In panels B-H, the blue, red, and green stars indicate the representative values for non-dialysis controls (DT(-), [β2m]_T_ = 2 μg/mL (0.17 μM) and [ALB]_T_ = 45 mg/mL (680 μM)) and dialysis patients before maintenance dialysis (DT(+, Pre), [β2m]_T_ = 50 μg/mL (4.24 μM) and [ALB]_T_ = 35 mg/mL (529 μM)) and dialysis patients after maintenance dialysis (DT(+, Post), [β2m]_T_ = 20 μg/mL (1.69 μM) and [ALB]_T_ = 38 mg/mL (574 μM)), respectively. The black arrows show the change in [β2m]_T_ and [ALB]_T_ by maintenance dialysis treatment. (I) Fluctuations of concentrations of the β2m monomer and serum albumin, temporary risk, *TR*, and accumulated risk, *AR*, as a function of the time. (J) Estimation of the onset of DRA by the accumulation of three risk factors at different conditions.

The equilibrium constants of Schemes 1–3 (i.e., Dissociation of serum albumin-native β2m complex (*K*_D_^C^), folding/unfolding (*K*_F_), and amyloid elongation (*K*_P_)) are represented by equations [1]–[3]:

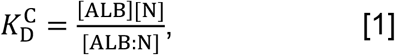

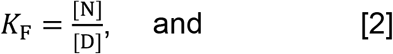

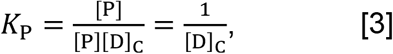

where [ALB:N], [ALB], [N], [D], and [P] refer to the molar concentrations of the serum albumin-native β2m complex, serum albumin, native β2m, denatured β2m, and amyloid fibrils, respectively, and [D]_C_ refers to the solubility of denatured β2m monomers. Although the seed-dependent elongation mechanism assumes the presence of preformed fibrils, *K*_P_ is independent of the concentration of seed fibrils. We considered that Scheme 3 and equation [3] are valid for amyloid fibril formation in general. [D]_C_ is also referred to as the “critical concentration” because amyloid fibrils form when the concentration of denatured monomers exceeds [D]_C_ ^17^.

#### Under supersaturation

We first obtained equilibrium concentrations of species under supersaturation where Schemes 1 and 2 are valid. With the total protein concentrations of serum albumin and β2m molecules as [ALB]_T_ and [β2m]_T_, respectively, [ALB:N], [N], and [D] are represented as functions of [ALB]_T_ and [β2m]_T_:

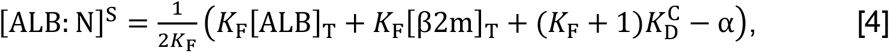

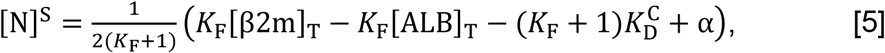

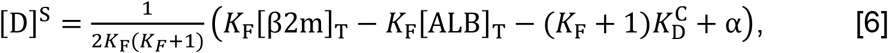

where 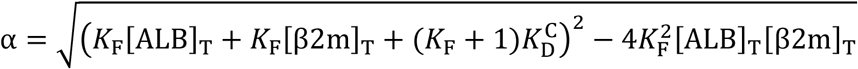. It should be noted that concentrations of the three species are linked by equations [1] and [2] and solving the quadratic equation is required to obtain the concentrations exactly. The 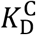 value was assumed to be 178 μM based on QCM analysis (Figure 5E) and the *K*_F_ value was considered to be 1650 based on a previous study ^46^.

Dependencies of respective species on total β2m and serum albumin concentrations in terms of concentrations ([ALB:N]^S^ and [D]^S^) or fractions (F_ALB:N_^S^ and F_N_^S^) are shown by red meshed surfaces in Figure 6B,C and Figure 6E,F, respectively. Figure 6C,F also shows [D]^S^ (= [N]^S^/*K*_F_) and F_D_^S^. Moreover, the degree of supersaturation (σ = [D]^S^/[D]_C_), which is a direct indicator of the risk of amyloid fibril formation and is proportional to [D]^S^, is shown in Figure 6C. The average values for non-dialysis controls and dialysis patients immediately before and after the maintenance dialysis treatment are indicated by blue, red, and green stars, respectively.

Upon increasing [β2m]_T_ in the absence of [ALB]_T_, the *σ* value increased (Figure 6C,F). However, this enhanced risk could be suppressed markedly by reducing D upon interaction with serum albumin. In other words, serum albumin works as a reservoir to reduce the risk of amyloid formation in terms of σ, or the law of mass action shifts the equilibrium to reduce D. Here, the [D]_C_ value was estimated to be 39.7 pM (SI Result 7), similar to the β2m concentration in healthy individuals and much lower than that in dialysis patients. However, no amyloid fibril was formed because of the high free energy barrier of supersaturation (Figure 6D,G).

Although we considered the simplest case whereby interaction occurs with the native β2m monomers, it is possible that denatured β2m monomers are also adsorbed by serum albumin, further decreasing the risk. On the other hand, the interactions between serum albumin and β2m monomer in vivo are likely to be weaker than observed here in an isolated in vitro system because of varying competing components in a serum milieu.

#### After breakdown of supersaturation

Then, under the conditions where [D]^S^ > [D]^C^, the new equilibrium establishes after breakdown of supersaturation by combining Schemes 1–3:

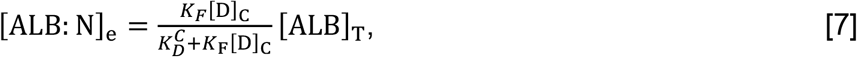

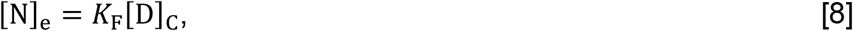

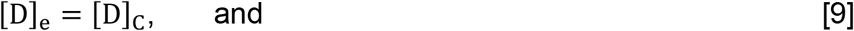

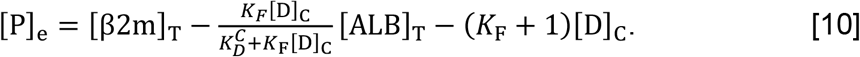

Most importantly, [D]_C_ is the dominant parameter determining the overall equilibrium; no consideration of a quadratic equation is necessary. Dependencies of respective species on [β2m]_T_ and [ALB]_T_ in terms of concentrations ([ALB:N]_e_, [N]_e_ and [P]_e_) or fractions (F_(ALB:N)e_ and F_(N)e_, and F_(P)e_) are shown by blue meshed surfaces in Figure 6B-D and Figure 6E-G, respectively. Figure 6C,F also shows [D]_e_ (= [N]_e_/*K*_F_) and F_(N)e_. Figure 6C,F shows the degree of supersaturation (σ), being 1 when [D]^S^ > [D]_C_ after breakdown of supersaturation, independent of β2m and serum albumin total concentrations. Except for the serum albumin-bound and native β2m species determined by [D]_C_, β2m monomers are converted to amyloid fibrils. The profiles suggest the marked impact of supersaturation on preventing amyloid fibril formation. The change in the concentrations of each β2m species is summarized in Supplementary Movie 1.

### Implication for supersaturation-limited onset of DRA

#### Roles of risk factors

Primary nucleation would be a rate-limiting step of amyloid fibril formation for hemodialysis patients ^47^. Once nuclei are formed, amyloid fibrils propagate rapidly throughout the patient’s body accelerated by secondary nucleation (i.e., fragmentations and lateral bindings), although it may take years until the onset of DRA. Thus, the potential for nucleation correlates with the risk of DRA onset. Based on the classical nucleation theory ^48^, the time for nucleation (i.e., lag time) shows an inverse correlation with the degree of supersaturation, σ, as ln(lag time) ∝ ln^-2^ *σ* (details in SI Note). Using the equation [6], we investigated the relative change in the lag time as a function of [β2m]_T_ and [ALB]_T_ (Figure 6H). As anticipated for dialysis patients (i.e., DT(+, Pre)), the higher β2m and lower serum albumin concentrations resulted in a shorter lag time.

During maintenance dialysis treatments, the relatively small β2m monomers (MW: ~11.8 kDa) pass through the pores of the dialysis membrane more than the larger serum albumin (MW: ~66.2 kDa). In parallel, maintenance dialysis removes a large amount of water from the patient’s body, concentrating the serum components. Combined effects are a decrease and an increase in β2m and serum albumin concentrations, respectively (Figure 6I). Consequently, the maintenance dialysis treatment reduces temporary risk (*TR*) for β2m amyloid fibril formation in a short term. However, the temporarily lowered risk gradually increases towards the next maintenance dialysis treatment: The β2m monomers newly produced accumulate and the concentration of serum albumin gradually decreases due to serum volume expansion by water uptake.

#### Onset of DRA

Considering the aforementioned fluctuation of *TR* for β2m amyloid fibril formation, we estimated the combined risk of DRA onset (Details in SI Note). First, the degree of supersaturation at each time-point, σ(t), was calculated as a function of β2m and serum albumin concentrations, which fluctuate during maintenance dialysis intervals (Figure 6I). Second, from the σ(t) value, the temporary risk, *TR*(*t*), of amyloid fibril formation was calculated as a function of time assuming that it is proportional to the inverse of the lag time, *TR*(*t*) ∝ ln^2^*σ*(*t*) (Figure 6I). Finally, the *TR*(*t*) value was integrated over the whole dialysis vintage to calculate the accumulated risk, *AR*(*t*), as 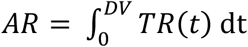, where DV denotes the dialysis vintage in years (Figure 6I).

Based on this model, we reproduced the onset of DRA (Figure 6J) as reported in the cohort study conducted by Hoshino *et al*. ^6^. For dialysis patients who developed DRA (*N* = 2157), the average dialysis vintage, β2m concentrations before and after maintenance dialysis treatments, and the serum albumin concentration before maintenance dialysis treatment were 18.2 years, [β2m]_T_^Pre^ =27.3 μg/mL (2.31 μM), [β2m]_T_^Post^ =10.2 μg/mL (0.86 μM), and [ALB]_T_^Pre^ = 36.6 mg/mL (553 μM), respectively. Assuming that the maintenance dialysis increases the concentration of serum albumin to 3 mg/mL (i.e., 8%) by condensation (i.e., [ALB]_T_^Post^=39.6 mg/mL (598 μM)), the *AR* value would reach ~2 at 18.2 years (black line in Figure 6J).

According to this mechanism, the reason why the patients with increased β2m concentrations (primary risk) and a long dialysis vintage (secondary risk) do not necessarily develop DRA ^4–8^ is explained by the variation in *AR* among individual patients. If the improvement of dialysis technology could reduce [β2m]_T_^Pre^ and [β2m]_T_^Post^ to 20 μg/mL (1.69 μM) and 5 μg/mL (0.42 μM), respectively, the onset of DRA would be delayed to 30.7 years (blue line in Figure 6J), markedly contributing to improvement of the quality of life of dialysis patients. Meanwhile, if the serum albumin level was kept at a healthy level (e.g., [ALB]_T_^Pre^ = 42 mg/mL (634 μM) and [ALB]_T_^Post^ = 45 mg/mL (680 μM)), the onset of DRA would be delayed to 20.3 years (red line in Figure 6J). The interaction of serum albumins with denatured β2m monomers, which we did not consider in this study, would further delay the onset of DRA. These considerations indicate that, although a decrease in the serum β2m concentration is the most effective way to prevent DRA, keeping the serum albumin concentration at a healthy level also contributes to the prevention of DRA by raising the inhibitory effects of the serum milieu on β2m amyloid formation.

#### Proteostasis network to stabilize supersaturation

To understand the pathophysiology of amyloidosis, it is essential to study the effects of aberrant mutations ^49,50^ and post-translational modifications ^51^ at the single molecule level. Furthermore, atomic-level observation of amyloid fibrils by cryo-electron microscopy ^52,53^ is also important to understand the relationship between polymorphisms in amyloid fibrils and the phenotype of amyloidosis. On the other hand, elucidating the reactive milieu of amyloid fibril formation from a physicochemical point of view gives us a novel perspective on how to reduce the risk of developing amyloidosis. In general, maintaining a healthy proteostasis network around amyloidogenic proteins under supersaturation may contribute to the prevention of not only DRA but also other amyloidoses. Although we succeeded in identifying a new risk factor that regulates β2m monomer supersaturation in the pathogenesis of DRA, the direct mechanisms that disrupt the supersaturation barrier are still unknown and their clarification will be essential for the prevention of DRA and other amyloidoses.

## Conclusions

Using sera collected from dialysis patients, we found that serum components inhibited β2m amyloid fibril formation in a concentration-dependent manner. We also revealed that, although the inhibitory effects were deteriorated in long-term dialysis patients, the deteriorated inhibitory effects were ameliorated by the maintenance dialysis treatment in the short term. Among serum components, we found that serum albumin plays a major role in maintaining the serum proteostasis network and thus supersaturation of β2m monomer via weak binding. Combining the elucidated risk factors (i.e., elevated β2m concentration, long dialysis vintage, and reduced serum albumin concentration), we constructed a model of developing DRA. The model indicates the importance of monitoring temporary risk (*TR*) and accumulated risk (*AR*) by a methodology like HANABI-2000, enabling a supersaturation-aided strategy to prevent the development of amyloidosis in general.

## Acknowledgement

The authors would like to thank Ph. D. Masatomo So for discussion. This study was performed as part of the Cooperative Research Program for the Institute for the Protein Research, Osaka University (CR-20-02) and was supported by the Japan Society for the Promotion of Science (20K22484), MDD Grand 2020 by the global center for medical engineering and informatics of Osaka University, Core-to-Core Program A (Advance Research Networks), Ministry of Education, Culture, Sports, Science and Technology (17H06352), and SENTAN from AMED (16809242).

## References

1 Chiti, F. & Dobson, C. M. Protein Misfolding, Amyloid Formation, and Human Disease: A Summary of Progress Over the Last Decade. Annu Rev Biochem 86, 27–68, doi:10.1146/annurev-biochem-061516-045115 (2017).

2 Benson, M. D. et al. Amyloid nomenclature 2020: update and recommendations by the International Society of Amyloidosis (ISA) nomenclature committee. Amyloid 27, 217–222, doi:10.1080/13506129.2020.1835263 (2020).

3 Gejyo, F. et al. A new form of amyloid protein associated with chronic hemodialysis was identified as beta 2-microglobulin. Biochem Biophys Res Commun 129, 701–706, doi:10.1016/0006-291x(85)91948-5 (1985).

4 Gejyo, F., Homma, N., Suzuki, Y. & Arakawa, M. Serum levels of beta 2-microglobulin as a new form of amyloid protein in patients undergoing long-term hemodialysis. N Engl J Med 314, 585–586, doi:10.1056/NEJM198602273140920 (1986).

5 Yamamoto, S. & Gejyo, F. Historical background and clinical treatment of dialysis-related amyloidosis. Biochim Biophys Acta 1753, 4–10, doi:10.1016/j.bbapap.2005.09.006 (2005).

6 Hoshino, J. et al. Significance of the decreased risk of dialysis-related amyloidosis now proven by results from Japanese nationwide surveys in 1998 and 2010. Nephrol Dial Transplant 31, 595–602, doi:10.1093/ndt/gfv276 (2016).

7 Scarpioni, R. et al. Dialysis-related amyloidosis: challenges and solutions. Int J Nephrol Renovasc Dis 9, 319–328, doi:10.2147/IJNRD.S84784 (2016).

8 Hatano, M. et al. Dialysis-related carpal tunnel syndrome in the past 40 years. Clin Exp Nephrol, doi:10.1007/s10157-021-02122-8 (2021).

9 Vincent, C. & Revillard, J. P. Beta-2-microglobulin and HLA-related glycoproteins in human urine and serum. Contrib Nephrol 26, 66–88, doi:10.1159/000396105 (1981).

10 Ohhashi, Y., Kihara, M., Naiki, H. & Goto, Y. Ultrasonication-induced amyloid fibril formation of beta2-microglobulin. J. Biol. Chem. 280, 32843–32848 (2005).

11 Yoshimura, Y. et al. Distinguishing crystal-like amyloid fibrils and glass-like amorphous aggregates from their kinetics of formation. Proc. Natl. Acad. Sci. U. S. A. 109, 14446–14451, doi:DOI 10.1073/pnas.1208228109 (2012).

12 Adachi, M., So, M., Sakurai, K., Kardos, J. & Goto, Y. Supersaturation-limited and Unlimited Phase Transitions Compete to Produce the Pathway Complexity in Amyloid Fibrillation. J Biol Chem 290, 18134–18145, doi:M115.648139/10.1074/jbc.M115.648139 (2015).

13 So, M., Hall, D. & Goto, Y. Revisiting supersaturation as a factor determining amyloid fibrillation. Curr Opin Struct Biol 36, 32–39, doi:S0959-440X(15)00174-8/10.1016/j.sbi.2015.11.009 (2016).

14 Noji, M. et al. Heating during agitation of beta2-microglobulin reveals that supersaturation breakdown is required for amyloid fibril formation at neutral pH. J Biol Chem 294, 15826–15835, doi:10.1074/jbc.RA119.009971 (2019).

15 Noji, M. et al. Breakdown of supersaturation barrier links protein folding to amyloid formation. Commun Biol 4, 120, doi:10.1038/s42003-020-01641-6 (2021).

16 Jarrett, J. T. & Lansbury, P. T., Jr. Seeding “one-dimensional crystallization” of amyloid: a pathogenic mechanism in Alzheimer’s disease and scrapie? Cell 73, 1055–1058, doi:0092-8674(93)90635-4 [pii] (1993).

17 Wetzel, R. Kinetics and thermodynamics of amyloid fibril assembly. Acc. Chem. Res. 39, 671–679, doi:10.1021/ar050069h (2006).

18 Lansbury, P. T., Jr. & Caughey, B. The chemistry of scrapie infection: implications of the ‘ice 9’ metaphor. Chem Biol 2, 1–5, doi:1074-5521(95)90074-8 (1995).

19 Coquerel, G. Crystallization of molecular systems from solution: phase diagrams, supersaturation and other basic concepts. Chemical Society Reviews 43, 2286–2300, doi:10.1039/c3cs60359h (2014).

20 Matsushita, Y. et al. Nanoscale Dynamics of Protein Assembly Networks in Supersaturated Solutions. Sci Rep 7, 13883, doi:10.1038/s41598-017-14022-7 (2017).

21 Wallace, A. F. et al. Microscopic evidence for liquid-liquid separation in supersaturated CaCO3 solutions. Science 341, 885–889, doi:341/6148/885/10.1126/science.1230915 (2013).

22 Zhang, C. M. et al. Possible mechanisms of polyphosphate-induced amyloid fibril formation of beta2-microglobulin. Proc Natl Acad Sci U S A 116, 12833–12838, doi:10.1073/pnas.1819813116 (2019).

23 Minton, A. P. Implications of macromolecular crowding for protein assembly. Curr Opin Struct Biol 10, 34–39, doi:10.1016/s0959-440x(99)00045-7 (2000).

24 Minton, A. P. The influence of macromolecular crowding and macromolecular confinement on biochemical reactions in physiological media. J Biol Chem 276, 10577–10580, doi:10.1074/jbc.R100005200 (2001).

25 Uversky, V. N., E, M. C., Bower, K. S., Li, J. & Fink, A. L. Accelerated alpha-synuclein fibrillation in crowded milieu. FEBS Lett 515, 99–103, doi:10.1016/s0014-5793(02)02446-8 (2002).

26 Seeliger, J., Werkmuller, A. & Winter, R. Macromolecular crowding as a suppressor of human IAPP fibril formation and cytotoxicity. PLoS One 8, e69652, doi:10.1371/journal.pone.0069652 (2013).

27 Nakajima, K. et al. Optimized sonoreactor for accelerative amyloid-fibril assays through enhancement of primary nucleation and fragmentation. Ultrason Sonochem 73, 105508, doi:10.1016/j.ultsonch.2021.105508 (2021).

28 Nakajima, K. et al. Half-Time Heat Map Reveals Ultrasonic Effects on Morphology and Kinetics of Amyloidogenic Aggregation Reaction. ACS Chem Neurosci 12, 3456–3466, doi:10.1021/acschemneuro.1c00461 (2021).

29 Goto, Y. et al. Development of HANABI, an ultrasonication-forced amyloid fibril inducer. Neurochem Int 153, 105270, doi:10.1016/j.neuint.2021.105270 (2022).

30 Powers, E. T., Morimoto, R. I., Dillin, A., Kelly, J. W. & Balch, W. E. Biological and chemical approaches to diseases of proteostasis deficiency. Annu Rev Biochem 78, 959–991, doi:10.1146/annurev.biochem.052308.114844 (2009).

31 So, M. et al. Ultrasonication-dependent acceleration of amyloid fibril formation. J. Mol. Biol. 412, 568–577, doi:S0022-2836(11)00847-3/10.1016/j.jmb.2011.07.069 (2011).

32 Umemoto, A., Yagi, H., So, M. & Goto, Y. High-throughput analysis of the ultrasonication-forced amyloid fibrillation reveals the mechanism underlying the large fluctuation in the lag time. J. Biol. Chem. 289, 27290–27299 doi:M114.569814/10.1074/jbc.M114.569814 (2014).

33 Nakajima, K. et al. Nucleus factory on cavitation bubble for amyloid beta fibril. Sci Rep 6, 22015, doi:10.1038/srep22015 (2016).

34 Nakajima, K. et al. Drastic acceleration of fibrillation of insulin by transient cavitation bubble. Ultrason Sonochem 36, 206–211, doi:10.1016/j.ultsonch.2016.11.034 (2017).

35 Naiki, H. et al. Establishment of a kinetic model of dialysis-related amyloid fibril extension in vitro. Amyloid 4, 223–232 (1997).

36 Chatani, E. et al. Ultrasonication-dependent production and breakdown lead to minimum-sized amyloid fibrils. Proc Natl Acad Sci U S A 106, 11119–11124, doi:10.1073/pnas.0901422106 (2009).

37 Kaneko, S. & Yamagata, K. Hemodialysis-related amyloidosis: Is it still relevant? Semin Dial 31, 612–618, doi:10.1111/sdi.12720 (2018).

38 Parnetti, L. et al. CSF and blood biomarkers for Parkinson’s disease. Lancet Neurol 18, 573–586, doi:10.1016/S1474-4422(19)30024-9 (2019).

39 Iimura, O., Tabei, K., Nagashima, H. & Asano, Y. A study on regulating factors of plasma refilling during hemodialysis. Nephron 74, 19–25, doi:10.1159/000189276 (1996).

40 Ogi, H. Wireless-electrodeless quartz-crystal-microbalance biosensors for studying interactions among biomolecules: a review. Proc Jpn Acad Ser B Phys Biol Sci 89, 401–417, doi:10.2183/pjab.89.401 (2013).

41 Noi, K., Iwata, A., Kato, F. & Ogi, H. Ultrahigh-Frequency, Wireless MEMS QCM Biosensor for Direct, Label-Free Detection of Biomarkers in a Large Amount of Contaminants. Anal Chem 91, 9398–9402, doi:10.1021/acs.analchem.9b01414 (2019).

42 Ogi, H. et al. Concentration dependence of IgG-protein A affinity studied by wireless-electrodeless QCM. Biosens Bioelectron 22, 3238–3242, doi:10.1016/j.bios.2007.03.003 (2007).

43 Ogi, H. et al. Nonspecific-adsorption behavior of polyethylenglycol and bovine serum albumin studied by 55-MHz wireless-electrodeless quartz crystal microbalance. Biosens Bioelectron 24, 3148–3152, doi:10.1016/j.bios.2009.03.035 (2009).

44 Nakano, S., Miyoshi, D. & Sugimoto, N. Effects of molecular crowding on the structures, interactions, and functions of nucleic acids. Chem Rev 114, 2733–2758, doi:10.1021/cr400113m (2014).

45 Munishkina, L. A., Cooper, E. M., Uversky, V. N. & Fink, A. L. The effect of macromolecular crowding on protein aggregation and amyloid fibril formation. J Mol Recognit 17, 456–464, doi:10.1002/jmr.699 (2004).

46 Kardos, J., Yamamoto, K., Hasegawa, K., Naiki, H. & Goto, Y. Direct measurement of the thermodynamic parameters of amyloid formation by isothermal titration calorimetry. J Biol Chem 279, 55308–55314, doi:10.1074/jbc.M409677200 (2004).

47 Ghosh, P., Vaidya, A., Kumar, A. & Rangachari, V. Determination of critical nucleation number for a single nucleation amyloid-beta aggregation model. Math Biosci 273, 70–79, doi:10.1016/j.mbs.2015.12.004 (2016).

48 Auer, S. & Frenkel, D. Prediction of absolute crystal-nucleation rate in hard-sphere colloids. Nature 409, 1020–1023, doi:10.1038/35059035 (2001).

49 Adams, D., Koike, H., Slama, M. & Coelho, T. Hereditary transthyretin amyloidosis: a model of medical progress for a fatal disease. Nat Rev Neurol 15, 387–404, doi:10.1038/s41582-019-0210-4 (2019).

50 Mizuno, H. et al. Dialysis-related amyloidosis associated with a novel beta2-microglobulin variant. Amyloid 28, 42–49, doi:10.1080/13506129.2020.1813097 (2021).

51 Radamaker, L. et al. Role of mutations and post-translational modifications in systemic AL amyloidosis studied by cryo-EM. Nat Commun 12, 6434, doi:10.1038/s41467-021-26553-9 (2021).

52 Fitzpatrick, A. W. P. et al. Cryo-EM structures of tau filaments from Alzheimer’s disease. Nature 547, 185–190, doi:10.1038/nature23002 (2017).

53 Iadanza, M. G. et al. The structure of a beta2-microglobulin fibril suggests a molecular basis for its amyloid polymorphism. Nat Commun 9, 4517, doi:10.1038/s41467-018-06761-6 (2018).

