## Supplementary Information for "Macromolecular crowding and supersaturation protect hemodialysis patients from the onset of dialysis-related amyloidosis"

### Supplementary Text

#### Supplementary Results

##### SI Result 1: High performance reversed-phase chromatography of aggregates formed by ultrasonication

High performance reversed-phase chromatographic analysis was performed to confirm that the aggregates formed by ultrasonication consisted of intact monomers that had not been fragmented by ultrasonication. The dissolution of preformed aggregates with 90% (v/v) DMSO under acidic conditions was confirmed using thioflavin-T (ThT) dye by spectrofluorometer (F-7100, HITACHI), as shown in Figure S3A. The intact monomer without ultrasonication was prepared as a control. Figure S3B shows the chromatograms of each sample. The amyloid fibrils dissolved by 90% DMSO (acidic) showed the same elution peak as the intact monomer at 15.76 min. Enlarged chromatograms are shown in the main body in Figure 1C. The chromatogram of the dissolved amyloid fibrils showed a sub-peak, as presented in Figure S3C, but the area under the sub-peak curve was 1.8%, which was negligible compared with the area under the main peak. These results demonstrated that amyloid fibrils formed by ultrasonication consisted of intact monomers without any ultrasonic fragmentations or other damage during the ultrasonication-based assay.

##### SI Result 2: Supplementary TEM images

To confirm the reproducibility of the TEM images of the main body (Figure 1D-F), multiple images acquired with the same sample were included as supplementary data, as shown in Figure S4. As in the TEM images of the main body, the ThT-positive aggregates formed by ultrasonication showed a weak-contrast fibrillar morphology

(Figure S4A), and the aggregates seeded and elongated from the amyloid fibrils formed by ultrasonication showed a clear-contrast fibrous morphology (Figure S4B). The ThT-negative aggregates formed in the presence of 15% (v/v) serum were amorphous (Figure S4C).

**SI Result 3: Serum concentration of β2m monomer in each patient cohort**

Serum samples were collected from non-dialysis controls (DT(-), $N$ = 30) and dialysis patients (DT(+), $N$ = 58). In the group of dialysis patients, 28 patients donated two serum samples, one collected immediately before (DT(+, Pre)) and one immediately after (DT(+, Post)) a maintenance dialysis treatment. The remaining 30 patients donated their serum samples collected immediately before (DT(+, Pre)) a maintenance dialysis treatment. Thus, 116 serum samples were used in this study.

For all sera, the concentrations of β2m monomer were measured with enzyme linked immunosorbent assay (ELISA) using a commercially available reagent kit (KGE019, Human beta 2-Microglobulin Parameter Assay Kit, R&D Systems). The results of quantification for all sera are shown in Figure S5.

The serum concentrations of β2m monomer were clearly higher in the serum of dialysis patients than those of non-dialysis controls, as shown in Figure S5A. The concentrations in dialysis patients and non-dialysis controls were 23.9 ± 5.2 ($N$ = 30) and 1.3 ± 0.4 μg/mL ($N$ = 30), respectively, which were consistent with the literature [1,2]. Meanwhile, the serum concentrations of β2m monomer markedly decreased after maintenance dialysis treatments, as shown in Figure S5B. The pre- and post-dialysis β2m concentrations were 24.9 ± 5.3 ($N$ = 28) and 7.7 ± 4.4 μg/mL ($N$ = 28), respectively, which were also consistent with the literature [1,2]. When serum was added to the recombinant β2m solution at a concentration of 5% (v/v), the concentration of β2m monomer from sera was at most 2 μg/mL, being much less than in standard solutions (1.0 mg/mL) for the HANABI assay, and its effects on amyloid fibril formation would be negligible.

**SI Result 4: Relationship between ThT fluorescence intensity and the amount of amyloid fibrils in the presence of sera**

In amyloid fibril formation, the addition of preformed fibrils bypasses the primary nucleation and accelerates amyloid formation, the so-called seeding reaction [3]. In general, the reaction time for amyloid formation is shortened in a seed-concentration-dependent manner, i.e., the higher seed concentrations induce more rapid amyloid fibril formation [4]. Based on the concentration-dependent manner of the seeding reaction, we qualitatively evaluated the relationship between the ThT fluorescence intensity of amyloid fibrils formed at various serum concentrations and the net amount of amyloid fibrils.

As shown in Figure S6A,B, the lag time of amyloid formation depended on the seed concentration, indicating that the shorter the lag time, the greater the amount of preformed amyloid fibrils in the seed solution. Next, the seeds formed at serum concentrations of 0-10% (v/v) were added to the monomer solution, and the concentration of amyloid fibrils was evaluated by the seeding reaction, as shown in Figure S6C. As summarized in Figure S6D, the higher serum concentration led to a longer lag time in the seeding reaction. Figure S6E shows the relationship between ThT fluorescence intensity and the lag time in the seeding reaction. A shorter lag time indicates a higher amyloid fibril concentration in the sample solution, which means that the fibril concentration in the sample solution positively correlates with ThT fluorescence intensity of the sample solution.

**SI Result 5: Examination of significance of change in the effects of serum on amyloid formation before and after maintenance dialysis treatments**

To examine the change in the effect of serum on amyloid fibril formation before and after maintenance dialysis treatment, the ThT fluorescence assay with ultrasonication was performed using sera collected from 28 patients immediately before and after maintenance dialysis treatments. ThT kinetics are shown in Figure S8A, where red and blue lines denote the results with sera collected before and after the treatment, respectively. For each sample, ThT kinetics were measured using multiple independent solutions ($n \geq 4$). Using the data, the significance of any change in the lag time and ThT fluorescence intensity between sera collected before and after maintenance dialysis treatments was investigated by the unpaired *t*-test, as shown in Figure S8B,C, respectively. In 28 patients examined, 16 and 23 patients showed significantly slower kinetics (lag time) and less resultant amyloid fibrils (ThT fluorescence intensity) after the maintenance dialysis treatments, indicating that the inhibitory effects of serum on fibril formation improved after a single dialysis treatment in more than half of the dialysis patients (Table S1). In the main text, the results of the paired *t*-test for each average value are shown in Figure 4C,D.

**SI Result 6: Correlation between serum components and β2m fibril formation**

The correlation of each serum component with the amyloid formation was analyzed by means of the Pearson correlation coefficient. The concentrations of 27 serum components were measured for each serum, their concentrations were plotted on the horizontal axis, and the lag time and ThT fluorescence intensity were plotted on the vertical axes, shown in Figure S9. For each plot, the absolute value of the Pearson correlation coefficient, $|r|$, was calculated, and is shown in the inset of Figure S9. The top

10 components with the highest correlation coefficients are summarized in Tables S2 and S3. Among them, we focused on five serum components that showed correlation coefficients of 0.5 or higher for both the lag time and ThT fluorescence intensity: β2m monomer, blood urea nitrogen, creatinine, ureic acid, and serum albumin. Here, it should be noted that the final concentration of β2m monomers was negligible because it was less than 1% of the concentration of recombinant β2m monomers included in the standard solution used in the HANABI-2000 ultrasonic assays. On the other hand, blood urea nitrogen is an indicator of the blood concentration of urea. Urea is a classical denaturant of native proteins [5], shifting the equilibrium from native folded monomers to denatured unfolded monomers in a concentration-dependent manner. We previously studied the urea-induced unfolding of β2m monomers under a neutral condition [6]. The results showed that the urea concentration at the midpoint of denaturation was ~5 M. The serum concentration of blood urea nitrogen was less than 100 mg/dL (Figure S9, panel 25), which corresponds to a urea concentration of less than 36 mM. Such a low-concentration urea fails to affect the β2m monomer stability and its amyloid fibril formation. Therefore, since the high correlation between these two serum components and amyloidogenecity can be attributed to a spurious correlation, these two serum components were excluded as candidates for factors affecting amyloid formation.

Effects of three other serum components: serum albumin, ureic acid, and creatinine, on β2m amyloid fibril formation were investigated, as shown in Figure S10. For the experiments, serum albumin, ureic acid, and creatinine were added to the β2m monomer solution with a monomer concentration of 0.3 mg/mL including 20 mM sodium phosphate (pH 7.4), 500 mM NaCl, and 5 μM ThT dye. The results showed that serum albumin greatly markedly β2m fibril formation in a concentration-dependent manner (Figure S10D), while ureic acid (Figure S10H) and creatinine (Figure S10L) did not.

**SI Result 7: Determination of the solubility of denatured β2m monomers under neutral conditions**

The β2m monomer concentration in the supernatant of the solution after amyloid fibril formation was measured as described in Supplementary Materials and Methods. We confirmed that β2m monomers did not precipitate under the ultracentrifugation conditions (100,000x*g* for 1 h). The monomer concentration in the supernatant was determined to be 772 $\pm$ 389 μg/mL, which is the concentration of the native monomers ($[N]_e$) because the anti-β2m antibody was used for detection in ELISA measurements.

As previously reported [7,8], after the breakdown of supersaturation, amyloid fibrils are in equilibrium with monomers. In other words, when the solution reaches equilibrium after amyloid fibril formation, the concentration of the denatured monomers is the same as its solubility, $[D]_C$. Then, we calculated the concentration of the denatured monomers from the concentration of the folded monomers, which were obtained from ELISA measurements, using the equilibrium constant of the folding reaction, $K_F = 1.65\times10^3$ [9]. Then, the $[D]_C$ value ($= [N]_e/K_F$) was calculated to be 39.7 pM.

#### Supplementary Note

##### Estimation of the onset risk of DRA

According to the classical nucleation theory, the nucleation rate of a solute in a supersaturated solution, *J*, is denoted as:

$$J \propto \exp\left(-\frac{\Delta G^{\ddagger}}{RT}\right), \qquad [1]$$

where $\Delta G^{\ddagger}$, *R*, and *T* denote the activation free energy, gas constant, and absolute temperature, respectively. It is known that the $\Delta G^{\ddagger}$ value depends on the degree of supersaturation, $\sigma$, written as [10]:

$$\Delta G^{\ddagger} \propto \ln^{-2}(\sigma). \qquad [2]$$

In the amyloid fibril formation, the fibrils are formed by aggregation of the supersaturated denatured monomers. Thus, in this study, the $\sigma$ value is denoted as:

$$\sigma = \frac{[\mathrm{D}]^{\mathrm{S}}}{[\mathrm{D}]_{\mathrm{C}}}$$

$$= \frac{1}{2[\mathrm{D}]_{\mathrm{C}}K_{\mathrm{F}}(K_F+1)}\left(K_{\mathrm{F}}[\beta 2\mathrm{m}]_{\mathrm{T}} - K_{\mathrm{F}}[\mathrm{ALB}]_{\mathrm{T}} - (K_{\mathrm{F}}+1)K_{\mathrm{D}}^{\mathrm{C}} + \alpha\right), \quad [3]$$

where $\alpha = \sqrt{\left(K_{\mathrm{F}}[\mathrm{ALB}]_{\mathrm{T}} + K_{\mathrm{F}}[\beta 2\mathrm{m}]_{\mathrm{T}} + (K_{\mathrm{F}}+1)K_{\mathrm{D}}^{\mathrm{C}}\right)^2 - 4K_{\mathrm{F}}^2[\mathrm{ALB}]_{\mathrm{T}}[\beta 2\mathrm{m}]_{\mathrm{T}}}$. Given that the time for nucleation, $t_{\mathrm{nuc.}}$, is inversely proportional to the nucleation rate [11], the following equation is derived:

$$\ln(t_{\mathrm{nuc.}}) \propto \ln(J^{-1}) \propto \ln^{-2}(\sigma). \qquad (\sigma > 1) \qquad [4]$$

It should be noted that the nucleation reaction never occurs when $\sigma \leq 1$. Here, we replaced $t_{\mathrm{nuc.}}$ as a lag time for amyloid fibril formation because amyloid fibrils rapidly grow once nuclei form. These relations were used to estimate the onset risk of DRA.

The fluctuation of the temporary risk of amyloid fibril formation with time was calculated based on the change in the total β2m and albumin concentrations with time, as:

$$[\beta 2\mathrm{m}]_\mathrm{T}(t) = [\beta 2\mathrm{m}]_\mathrm{T}^\mathrm{Post} + \frac{[\beta 2\mathrm{m}]_\mathrm{T}^\mathrm{Pre} - [\beta 2\mathrm{m}]_\mathrm{T}^\mathrm{Post}}{T_D} t, \qquad [5]$$

$$[\mathrm{ALB}]_\mathrm{T}(t) = [\mathrm{ALB}]_\mathrm{T}^\mathrm{Post} + \frac{[\mathrm{ALB}]_\mathrm{T}^\mathrm{Post} - [\mathrm{ALB}]_\mathrm{T}^\mathrm{Pre}}{T_D} t, \qquad [6]$$

where $[\beta 2\mathrm{m}]_\mathrm{T}(t)$, $[\beta 2\mathrm{m}]_\mathrm{T}^\mathrm{Pre}$, and $[\beta 2\mathrm{m}]_\mathrm{T}^\mathrm{Post}$ are the total concentrations of β2m monomer at time *t* and immediately before and after maintenance dialysis treatment, respectively. The definition of super- and subscripts are the same as for the total concentration of serum albumin. $T_D$ is the interval between the maintenance dialysis treatments. By substituting equations [18] and [19] into equation [16], the degree of supersaturation at time $t$, $\sigma(t)$, was calculated, and, by a relation: $TR = \ln^2(\sigma(t))$, the temporary risk of amyloid fibril formation at time *t* was calculated.

#### Supplementary Materials and Methods

##### Recombinant β2m solutions

The recombinant β2m monomer with an additional methionine residue at the N terminus was expressed using *Escherichia coli* and purified as previously described [12]. The lyophilized β2m monomer was stored at -20 °C directly prior to experiments. The monomer was dissolved into deionized water and filtrated using a pore filter with a pore diameter of 220 nm. The monomer solution was mixed with other chemicals to their final concentrations as follows: [β2m] = 1.0 mg/mL; [NaPi(pH 7.4)] = 20 mM; [NaCl] = 300 mM; and [ThT] = 5 μM. In the amyloid formation experiments, the serum samples were added to the recombinant β2m solution at a volume ratio between 0.3-15% (v/v).

##### Collection and treatment of serum samples

**(i) Comparison between dialysis patients and non-dialysis controls.** We recruited 30 patients undergoing dialysis treatment and 30 non-dialysis controls in a single center.

In the dialysis patient group, dialysis treatments were 4-5 sessions conducted three times weekly with standard bicarbonate dialysate ($Na^+$: 140 mEq/L, $K^+$: 2.0 mEq/L, $Ca^{2+}$: 2.75 mEq/L, $Mg^{2+}$: 1.0 mEq/L, $Cl^-$: 112.25 mEq/L, and $HCO^{3-}$: 27.5 mEq/L) and dialyzers with synthetic polysulfone membranes. Baseline data, including: age, sex, body mass index, cause of kidney disease, systolic blood pressure, and serum levels of urea nitrogen, creatinine, serum albumin, blood hemoglobin, and C-reactive protein, were measured in both groups. Duration of dialysis treatment and single pool $Kt / V_{urea}$; were also reported in the dialysis patient group, and the estimated glomerular filtration rate (eGFR) was measured in the non-dialysis group. In the dialysis patient group, sera were collected before maintenance dialysis treatment. Continuous variables are expressed as medians (interquartile range). This study adhered to the Declaration of Helsinki and was approved by the Central Ethics Committee of Niigata University (2018-0054). All patients provided written informed consent.

Supplementary File 1A,B shows the demographic and clinical characteristics of the non-dialysis controls ($N$ = 30) and dialysis patients ($N$ = 30), demonstrating typical data for end-stage kidney disease.

**(ii) Investigation of the effects of a single maintenance dialysis treatment on the degree of inhibitory effects.** We recruited 28 patients undergoing long-term dialysis treatment in multi-centers. Dialysis treatments were 4-5 h sessions conducted three times weekly with standard bicarbonate dialysate ($Na^+$: 140 mEq/L, $K^+$: 2.0 mEq/L, $Ca^{2+}$: 2.75 mEq/L, $Mg^{2+}$: 1.0 mEq/L, $Cl^-$: 112.25 mEq/L, and $HCO^{3-}$: 27.5 mEq/L) and dialyzers with synthetic polysulfone membranes. Baseline data, including: age, sex, body mass index, cause of kidney disease, systolic blood pressure, and serum levels of serum albumin, blood hemoglobin, calcium, phosphorus, parathyroid hormone, β2m, and C-reactive protein, were measured in both groups. Duration of dialysis treatment and single pool $Kt / V_{urea}$; were also reported. The patients underwent long-term dialysis treatment. Sera were collected before and after maintenance dialysis treatment. Continuous variables are expressed as medians (interquartile range). This study adhered to the Declaration of Helsinki and was approved by the Central Ethics Committee of Niigata University (2018-0054). All patients provided written informed consent.

Supplementary File 1C,D shows the demographic and clinical characteristics of the long-term dialysis patients ($N$ = 28), demonstrating usual data as end-stage kidney disease.

**Ultrasonic assays of amyloid fibril formation using HANABI-2000**

We used an originally developed ultrasonication system, HANABI-2000, which is optimized for accelerated amyloid fibril formation [13]. The 198-μL sample solutions were added to a 96-well plate (675096, Greiner) and sealed with plastic film (547-KTS-HC, Watson). The sample solutions inside the plate were irradiated with ultrasound at a frequency of ~30 kHz, an optimized frequency for accelerating amyloid formation [14]. During the experiments, ultrasonication was performed with duty cycles comprising 0.3-s irradiation and 30-s quiescence incubation. To monitor the kinetics of amyloid fibril formation, the ThT fluorescence intensity was measured with excitation and emission wavelengths of 450 and 490 nm, respectively. Fluorescence measurements were performed every 10 min until the end of the experiment. The temperature of the sample solution was kept at 60 °C. Scheme of the HANABI assay is described in Figure S2.

**Circular dichroism (CD) spectrum measurements**

The secondary structure of the aggregates was analyzed using a CD spectrometer (JASCO Corp., J-820). After HANABI assays, the sample solutions had their β2m concentration adjusted to 0.15 mg/mL by dilution with deionized water. The

170-μL solution was injected into a quartz cell (JASCO Corp., 1103-0172) with a light path of 1 mm. The spectrum was acquired at a wavelength between 200 and 250 nm.

**High performance reversed-phase chromatography**

First, the fibril solution was diluted 10-fold with dimethyl sulfoxide (DMSO) and incubated for 1 h at room temperature [15]. Then, 6 M HCl solution was added to completely dissolve the aggregates. For the analysis, we used a high-performance liquid chromatography system (GILSON) with a C4 300-Å column (5C4-AR-300, COSMOSIL). The analysis was performed three times for each sample.

**Transmission electron microscopy (TEM) observation**

A 10-μL aliquot of the sample solution was placed on a collodion-coated copper grid (Nisshin EM Co.) for 1 min, and the remaining solution was removed with filter paper. The sample was then stained with a 1% (w/v) uranyl acetate solution for 1 min. Finally, the surface of the grid was rinsed with deionized water. TEM observation was performed using an H-7650 transmission electron microscope (HITACHI) with an acceleration voltage of 80 kV.

**Seeding experiments**

At first, the seed amyloid fibrils were prepared by ultrasonication at a β2m monomer concentration of 1.0 mg/mL under the conditions without sera. Then, the seeds were added to the monomer solutions ([β2m] = 0.1 mg/mL, [NaPi(pH7.4)] = 20 mM, [NaCl] = 300 mM, and [ThT] = 5 μM) at various seed concentrations. The amount of amyloid fibrils in aggregates prepared in the presence of sera was assayed by performing the same seeding reactions as above and comparing kinetics. The seeding experiments used intermittent shaking agitation with a cycle composed of 20-s shaking with revolution of 650 rpm and 580-s quiescent incubation. The temperature was 45 °C.

**Quartz crystal microbalance (QCM) measurements**

We investigated the interactions between β2m and serum albumin using a wireless-electrodeless quartz crystal microbalance (QCM) biosensor [16,17]. AT-cut quartz resonators with a fundamental frequency of 65 MHz and an in-plane size of 1.8×1.6 $mm^2$ were used. Films of 2-nm chromium and 15-nm gold were deposited on both sides of the quartz plate. QCMs were first cleaned using piranha solution (98% $H_2SO_4$: 30% $H_2O_2$ = 7:3) and deionized water. Then, 10 mM self-assembled monolayer (SAM) molecules in absolute ethanol were injected, with subsequent incubation overnight at 4 °C to make the linker layer. After washing with absolute ethanol and deionized water, a mixture of 100 mM *N*-hydroxysuccinimide and 100 mM 1-(3-dimethylaminopropyl)-3-ethylcarbodiimide hydrochloride in deionized water was injected, followed by incubation at room temperature for 2 h to activate the SAM termini, and then QCMs were rinsed with deionized water. Subsequently, 150 μM β2m monomer in buffer solution (20 mM NaPi, pH 7.4) was injected on both sides of QCM and incubated for 2 h at room temperature to immobilize the β2m monomer on the QCM surface. Finally, serum albumin or PEG solution with various concentrations was flowed using a micropump. The sensor cell was immersed in a water bath to keep the temperature at 37 °C during the measurement. The solution flow rate during measurement was ~0.5 mL/min.

**Solution Nuclear Magnetic Resonance (NMR) Measurements**

$^{15}$N-uniformly labelled β2m solutions at 1.0 mg/mL containing 20 mM sodium phosphate pH 7, 100 mM NaCl, 2% $D_2O$, and varying concentrations of serum albumin were prepared for 2D $^{1}$H-$^{15}$N heteronuclear single quantum coherence (HSQC) experiments. The spectra were recorded on a Bruker Avance III 500 MHz spectrometer with a cryogenic probe at 37 °C. The spectrometer was operated at $^{1}$H frequency of 500.13 MHz and $^{15}$N frequency of 50.68 MHz. $^{1}$H-$^{15}$N HSQC spectra were acquired with 2,048 complex points covering 7002.8 Hz for $^{1}$H and 192 complex points covering 1419.1 Hz for $^{15}$N. Resonance frequencies in the spectra were identified using the chemical shift lists on β2m [18]. NMR data were processed by TOPSPIN-NMR software and analyzed using Sparky [19].

**Solubility Measurements**

To address β2m amyloid fibril formation based on the supersaturation-limited mechanism, we measured the solubility of β2m monomer using an ultracentrifugation method combined with an ELISA assay [20,21]. First, β2m solution containing [β2m] = 1.0 mg/mL, [NaPi (pH7.4)] = 20 mM, [NaCl] = 300 mM, [ThT] = 5 μM was prepared, and then amyloid fibril formation was induced by ultrasonication using the HANABI-2000 instrument. After a sufficient reaction time over 20 h, the solution reached an equilibrium state with amyloid fibrils, which was confirmed by the saturation of ThT fluorescence intensity. The equilibrated solution was ultracentrifugated at 100,000*xg* for 1 h, and then, the concentration of β2m monomers in the supernatant was determined by the ELISA method.

**Supplementary Movie 1:** (i) TR under supersaturation: β2m species are in apparent equilibrium between three states, serum albumin-native monomer complex, native monomer, and denatured monomer due to the barrier of supersaturation. TR values vary with the total concentration of β2m and serum albumin. Maintenance dialysis treatment temporarily alleviates TR values by lowering the β2m concentration and increasing the serum albumin concentration. (ii) AR during long dialysis vintage: The risk for the onset of DRA accumulates during a long dialysis vintage. (iii) Breakdown of supersaturation and onset of DRA: When the AR value reaches a threshold, supersaturation is broken, and then, amyloid fibril formation begins. When [D] becomes $[D]_c$, amyloid formation finishes, and the entire solution is truly in equilibrium.

**Supplementary File 1:** (A) Demographic and clinical characteristics of the non-dialysis controls (*N* = 30) and dialysis patients (*N* = 30). (B) Summary of clinical data for the non-dialysis control and dialysis patient groups. (C) Demographic and clinical characteristics of the dialysis patients (*N* = 28) who were collected before and after a single maintenance dialysis treatment. (D) Summary of clinical data for the dialysis patients in sheet C.

#### Supplementary Figures

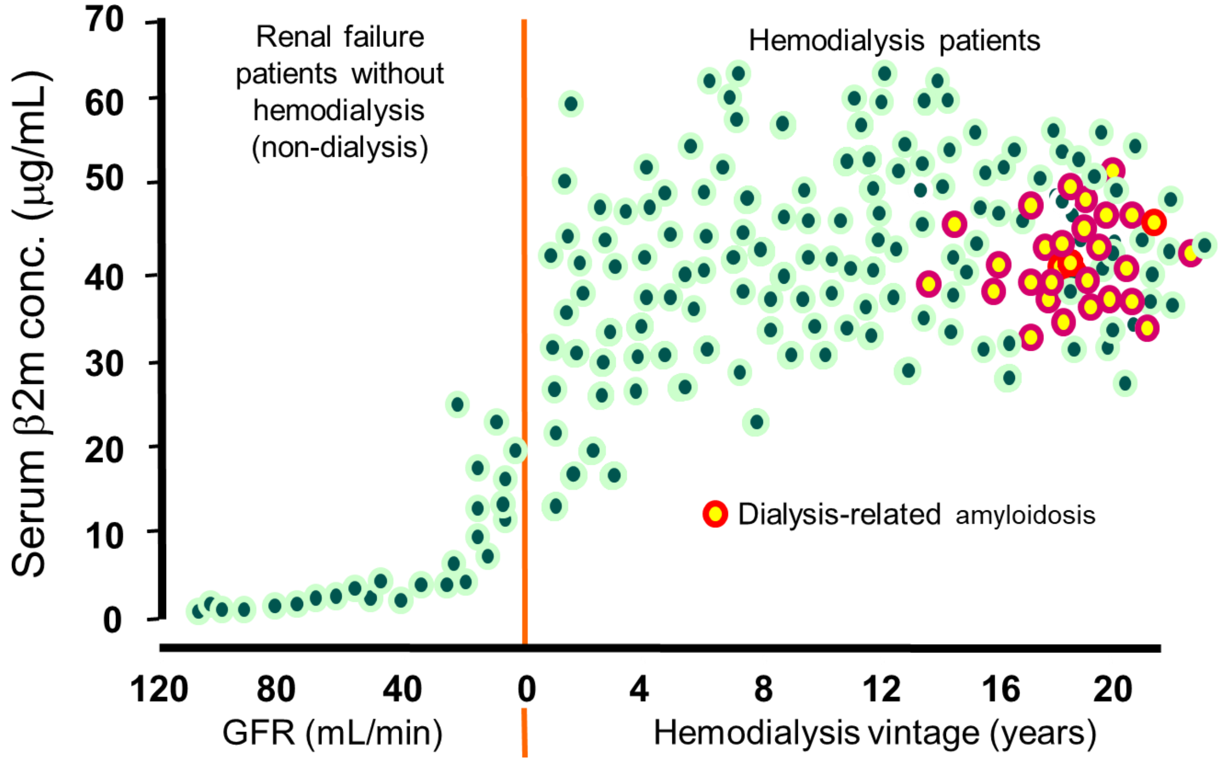


**Figure S1.** Serum β2m concentrations in chronic renal failure patients without hemodialysis vs. with hemodialysis [22]. There is an inverse correlation between the serum β2m concentration and glomerular filtration rate (GFR) in chronic renal failure patients. After the start of hemodialysis, the serum β2m concentration shows a rapid increase and becomes markedly elevated, exceeding the upper limit of normal by 40 times or more. The yellow dots indicate patients with dialysis-related amyloidosis, and the green dots represent patients without amyloidosis. The fact that no differences are apparent in the serum β2m concentrations between dialysis patients with and without amyloidosis indicates that, although the increase in the serum β2m concentration from a healthy level and long dialysis vintage are the primary and secondary risk factors for the onset of DRA, respectively, there are unknown risk factors controlling the onset (23).

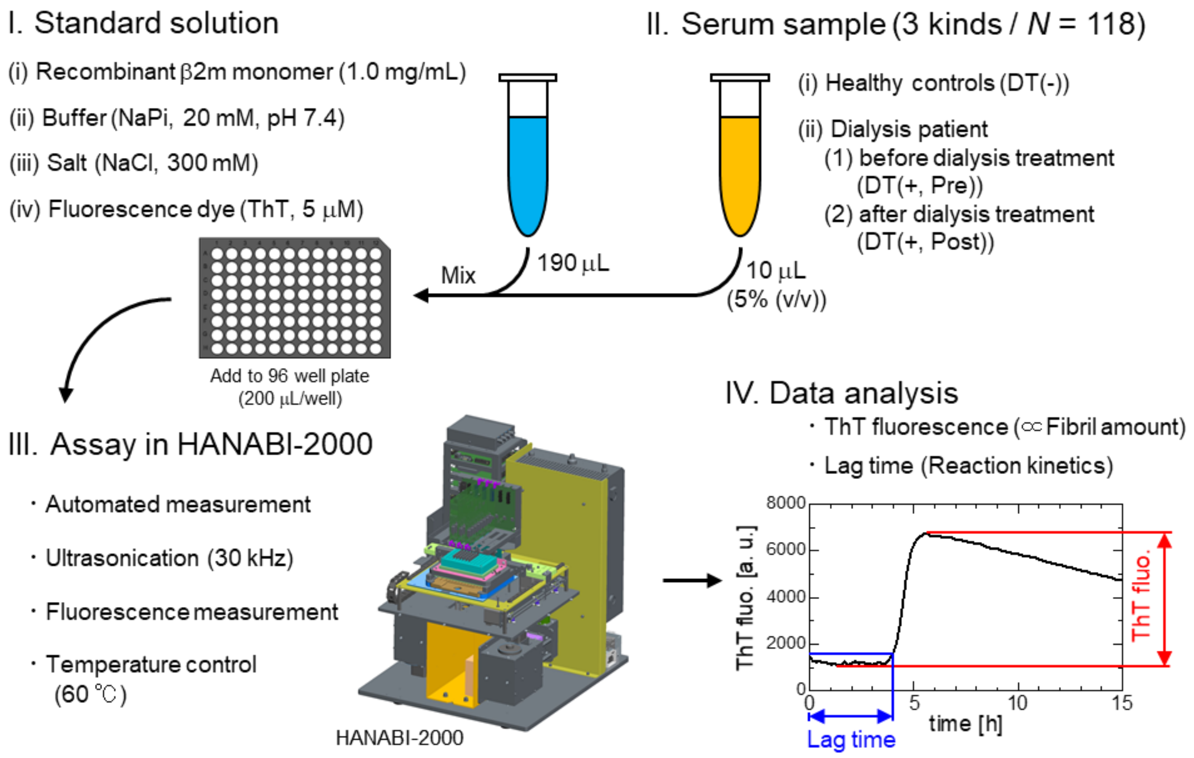


**Figure S2.** Schematic illustration of experimental scheme using HANABI-2000 instrument. To examine the effects of sera on the β2m amyloid fibril formation, the sera were added to the standard solution with a volume ratio of 5% (v/v). The aliquots were dispensed in a 96-well plate with the volume of 200 μL/well. The prepared plate was set to the HANABI-2000 instrument and was ultrasonicated at 60 °C. The fibril formation kinetics was monitored by ThT fluorescence measurement. The obtained ThT kinetics was analyzed in terms of its maximum intensity and lag time.

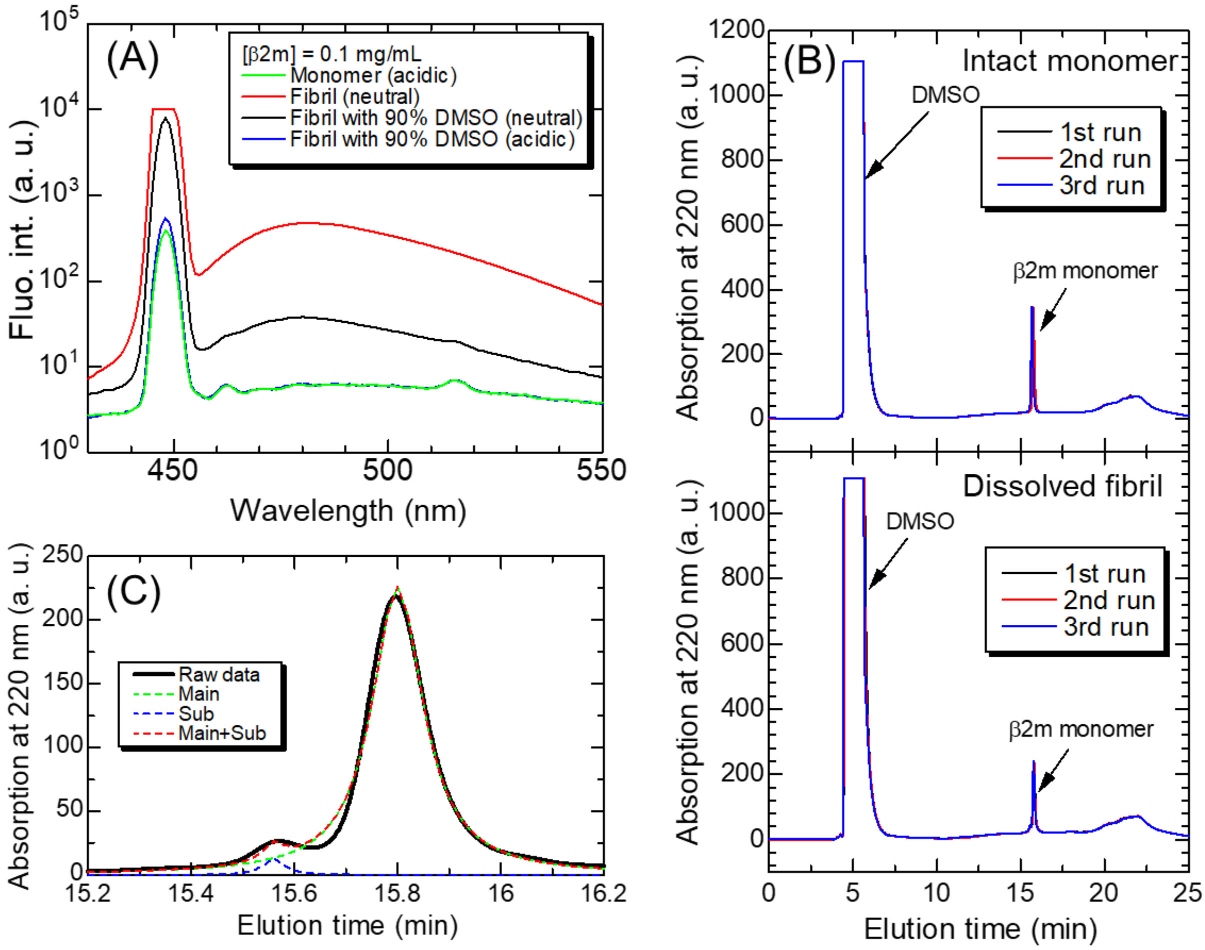


**Figure S3.** (A) Amyloid-specific ThT fluorescence spectra of intact monomers and depolymerized amyloid fibrils. Fluorescence measurements were performed with an excitation wavelength of 445 nm. (B) Reversed-phase chromatograms of the intact monomers (upper) and depolymerized amyloid fibrils (lower). The analysis was performed three times for each sample. (C) Enlarged view of the elution peak of the depolymerized amyloid fibrils (solid line) and fitted curves with the Lorentz function (dotted lines).

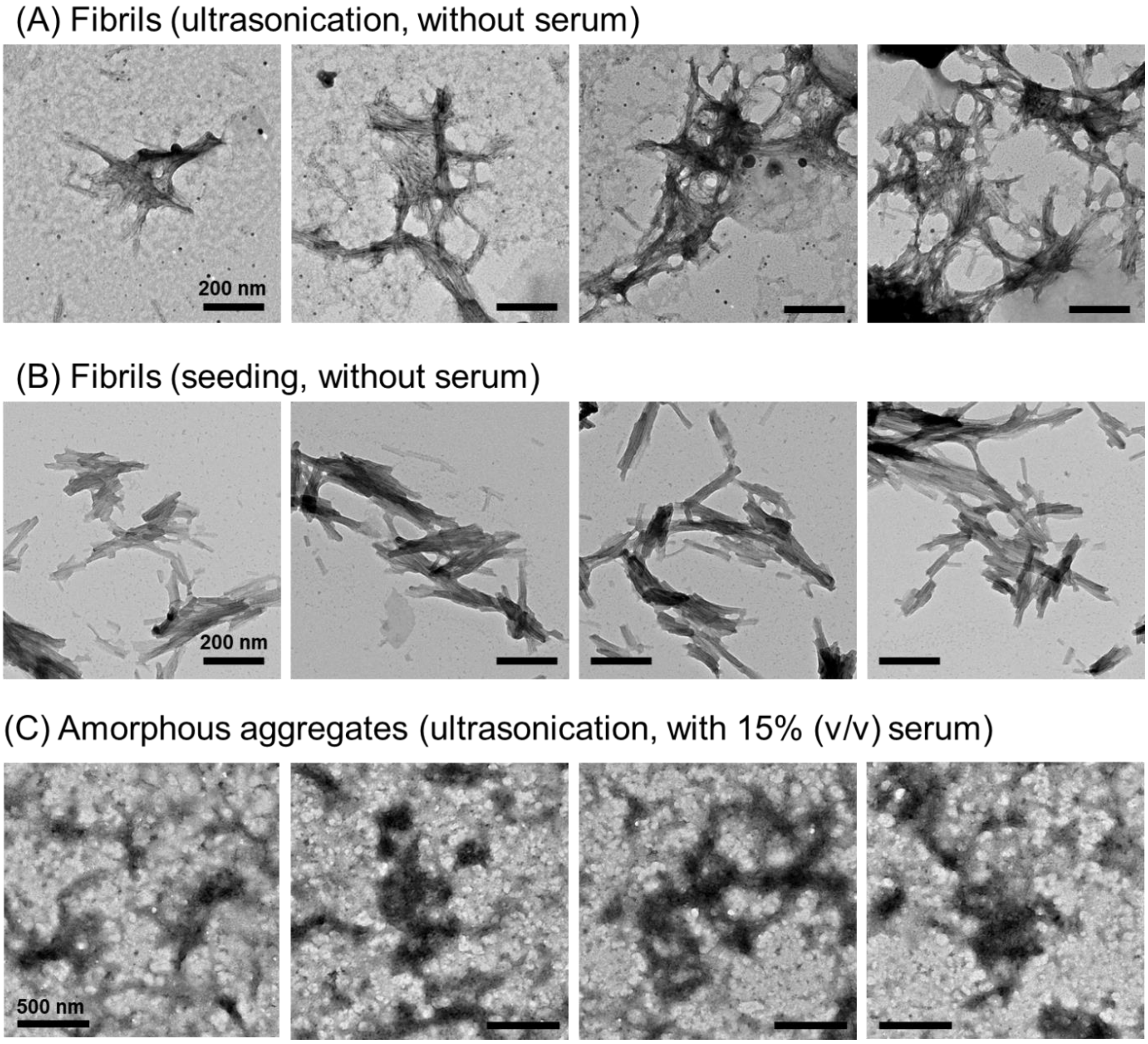


**Figure S4.** Supplementary TEM images of (A) amyloid fibrils formed by ultrasonication without serum addition, (B) amyloid fibrils seeded and elongated from the amyloid fibrils formed by ultrasonication, and (C) amorphous aggregates formed by ultrasonication with 15% (v/v) serum addition. Scale bars in panels A, B, and C denote 200, 200, and 500 nm, respectively.

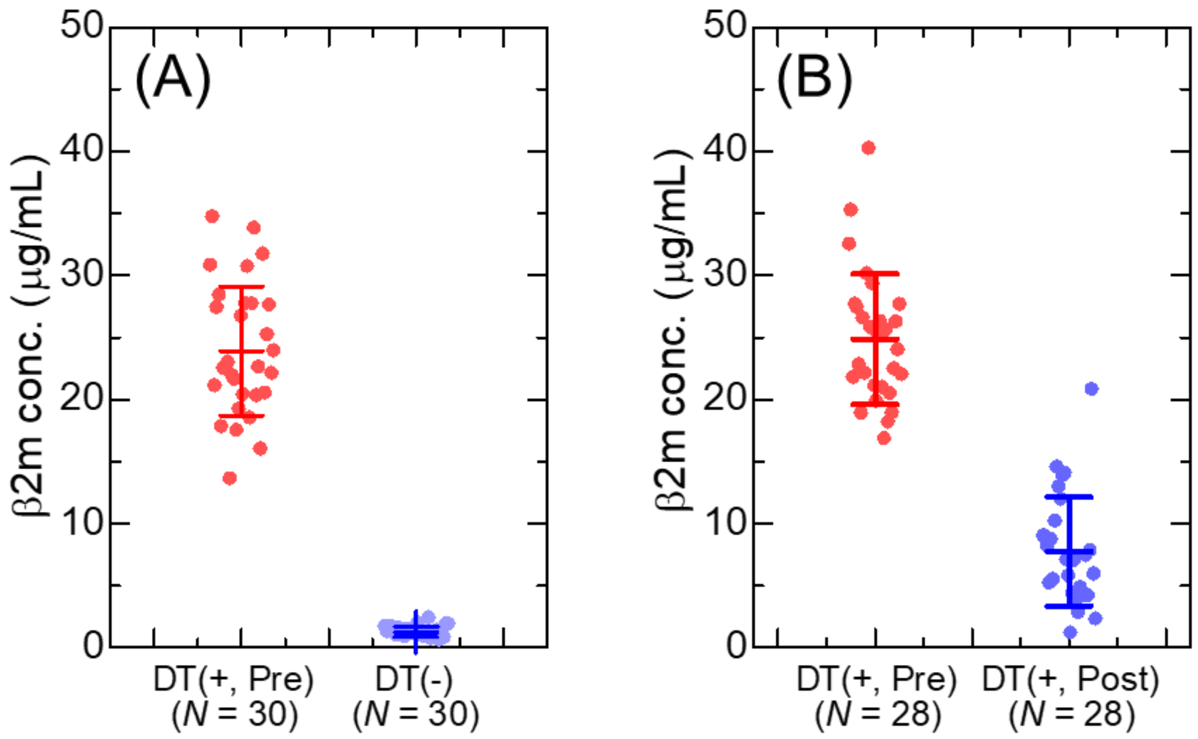


**Figure S5.** β2m monomer concentrations in sera used in a series of experiments shown in (A) Figure 3 and (B) Figure 4 of the main body. The sera of DT(+, Pre) in panels A and B were collected from different patient cohorts.

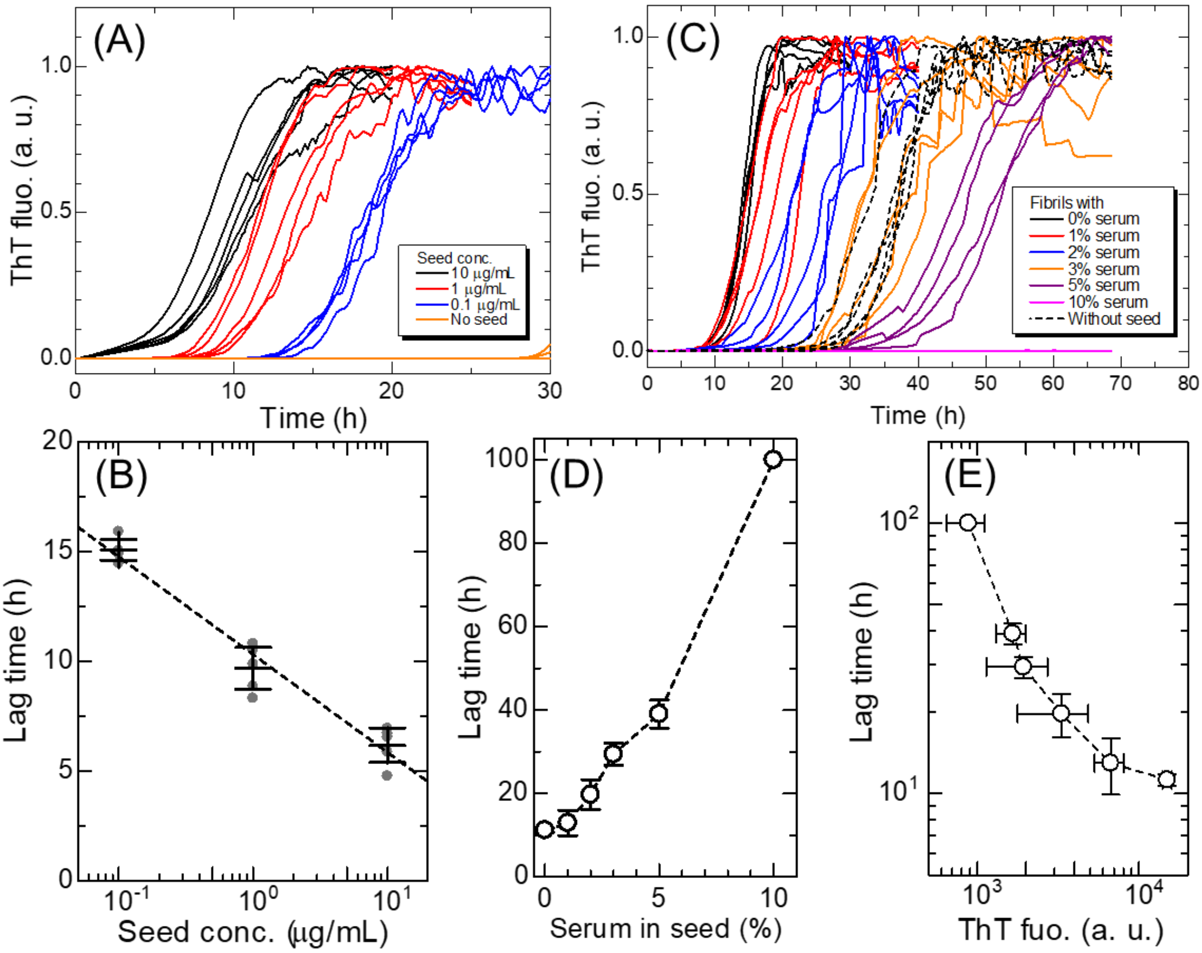


**Figure S6.** (A) ThT fluorescence kinetics of the seeding reaction at various seed concentrations (0.1, 1, and 10 μg/mL) and without seeds. (B) Relationship between the seed concentration and lag time in reactions obtained from the panel A. The error bars denote the standard deviation ($n$ = 5). (C) ThT fluorescence kinetics of the seeding reaction using seeds formed at various serum concentrations from 0-10% (v/v). Relationship between the lag time in the seeding reaction and (D) serum concentration in the seed solution and (E) ThT fluorescence intensity of the seed solution. The error bars denote the standard deviation ($n$ = 4).

(A)

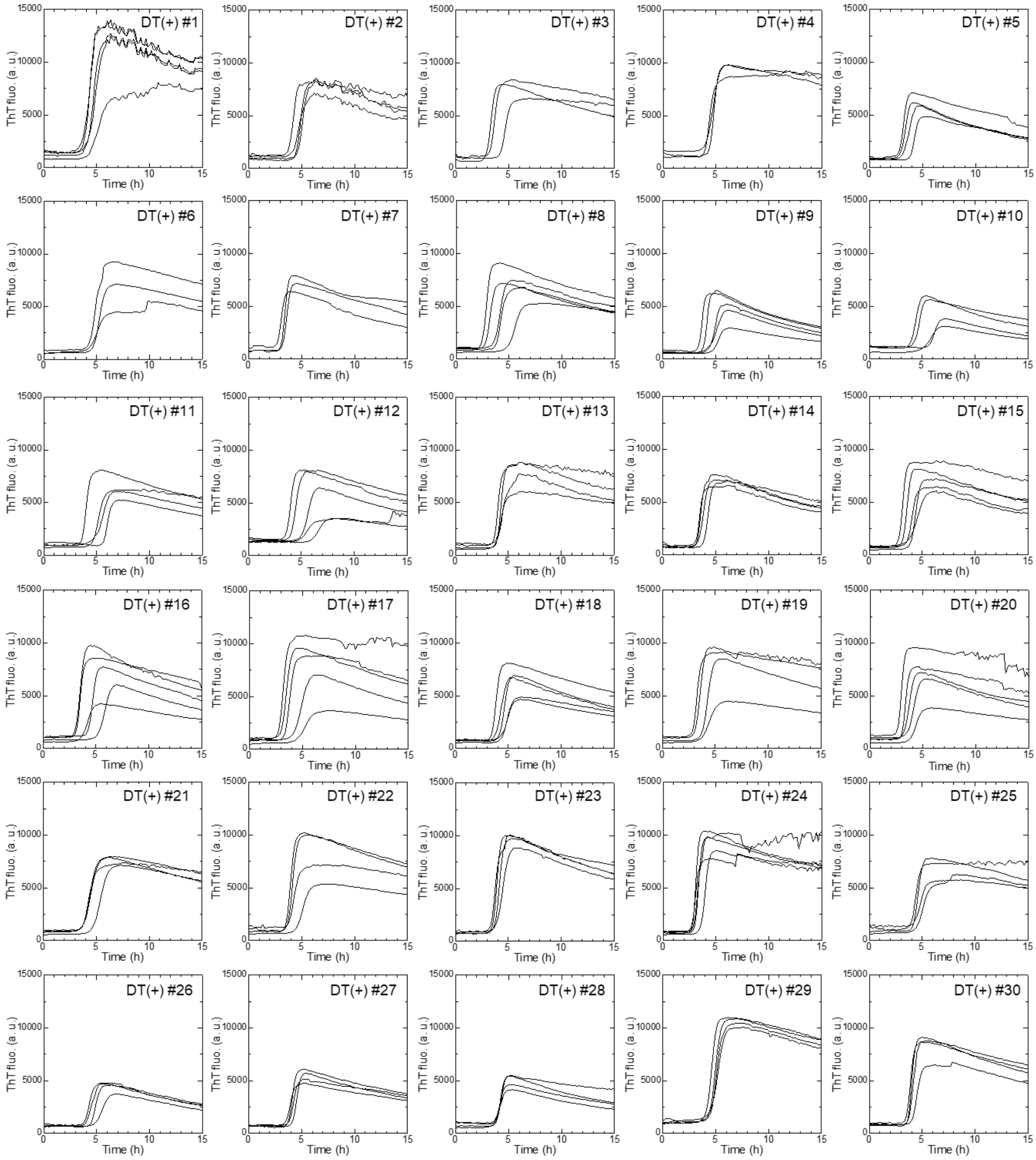


(Continue to the next page)

(B)

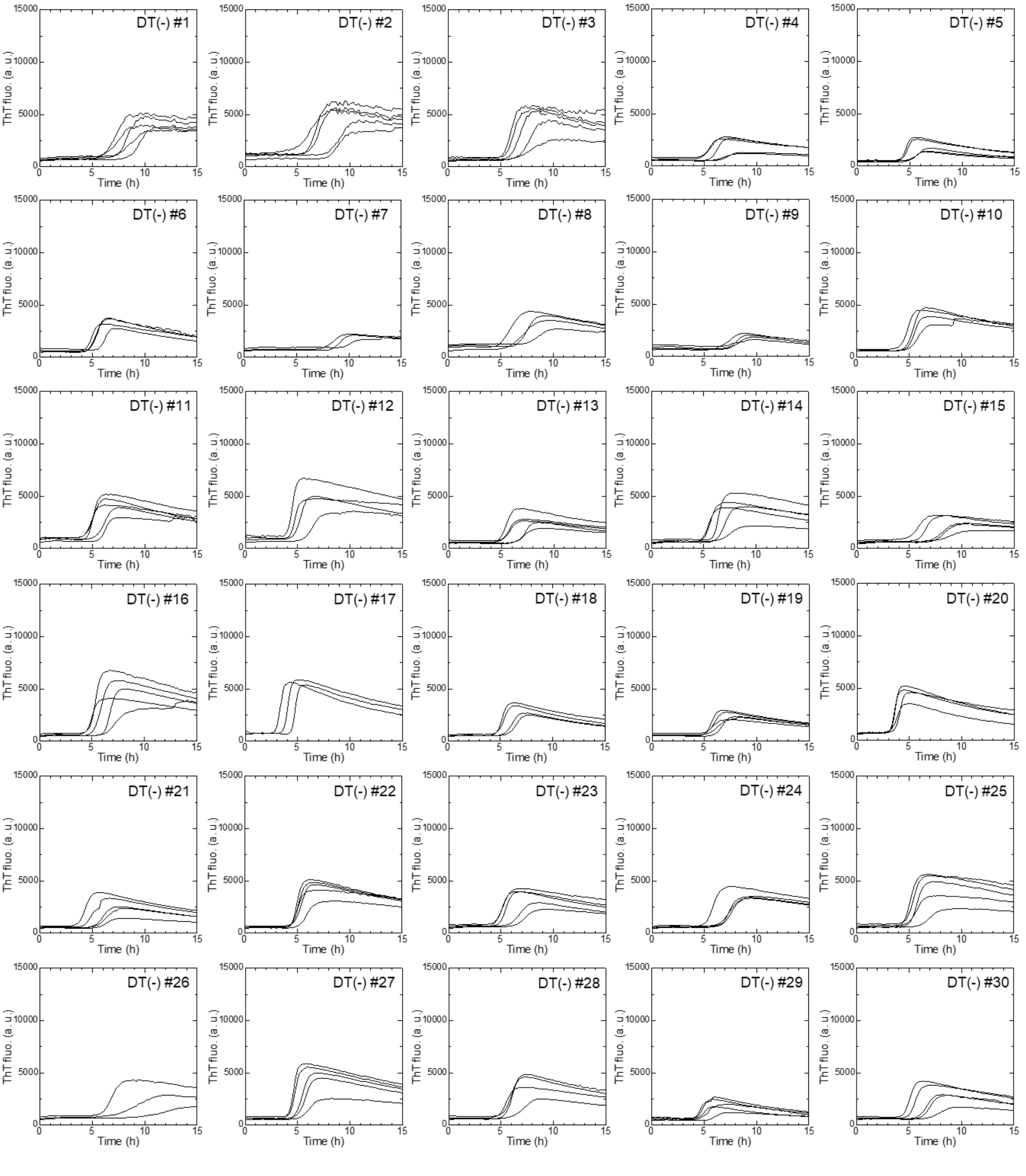


(Continue to the next page)

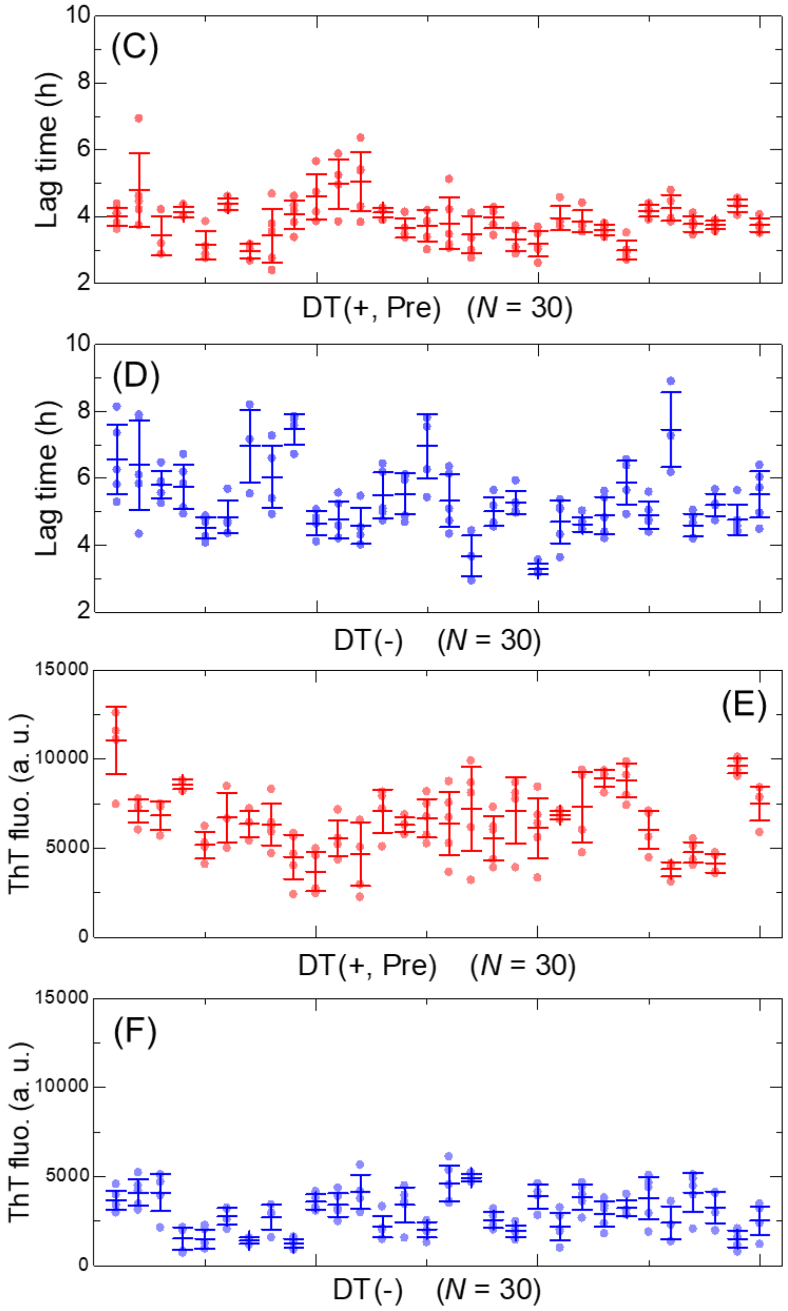


**Figure S7.** ThT kinetics of amyloid fibril formation using (A) sera from dialysis patients ($N$ = 30) and (B) sera from non-dialysis controls ($N$ = 30). For each serum, measurements were performed using at least three independent sample solutions ($n \geq 3$). (C-F) Summary of the ThT fluorescence assay for all samples in terms of the (C,D) lag time and (E,F) ThT fluorescence intensity, respectively. Error bars denote the standard deviation.

(A)

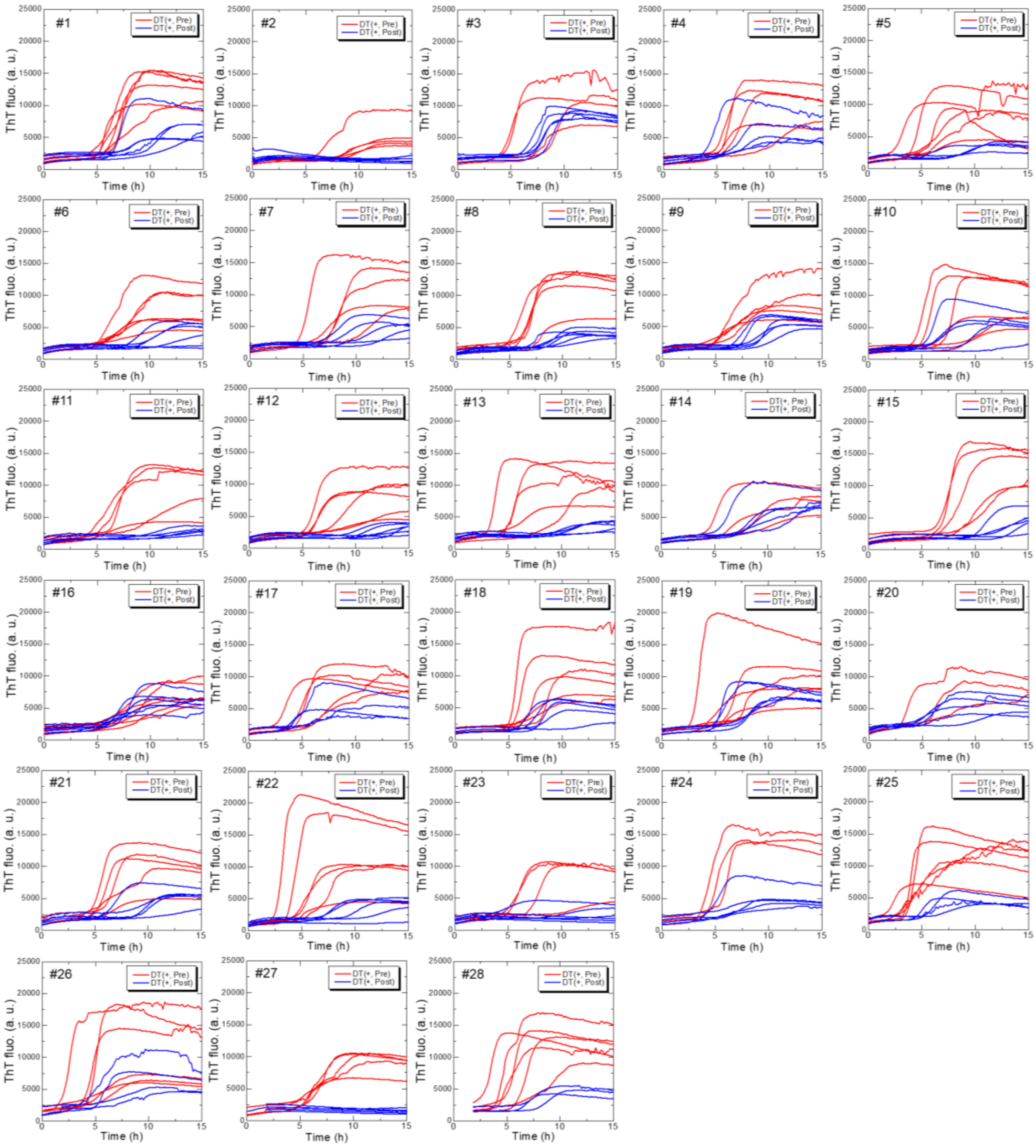


(Continue to the next page)

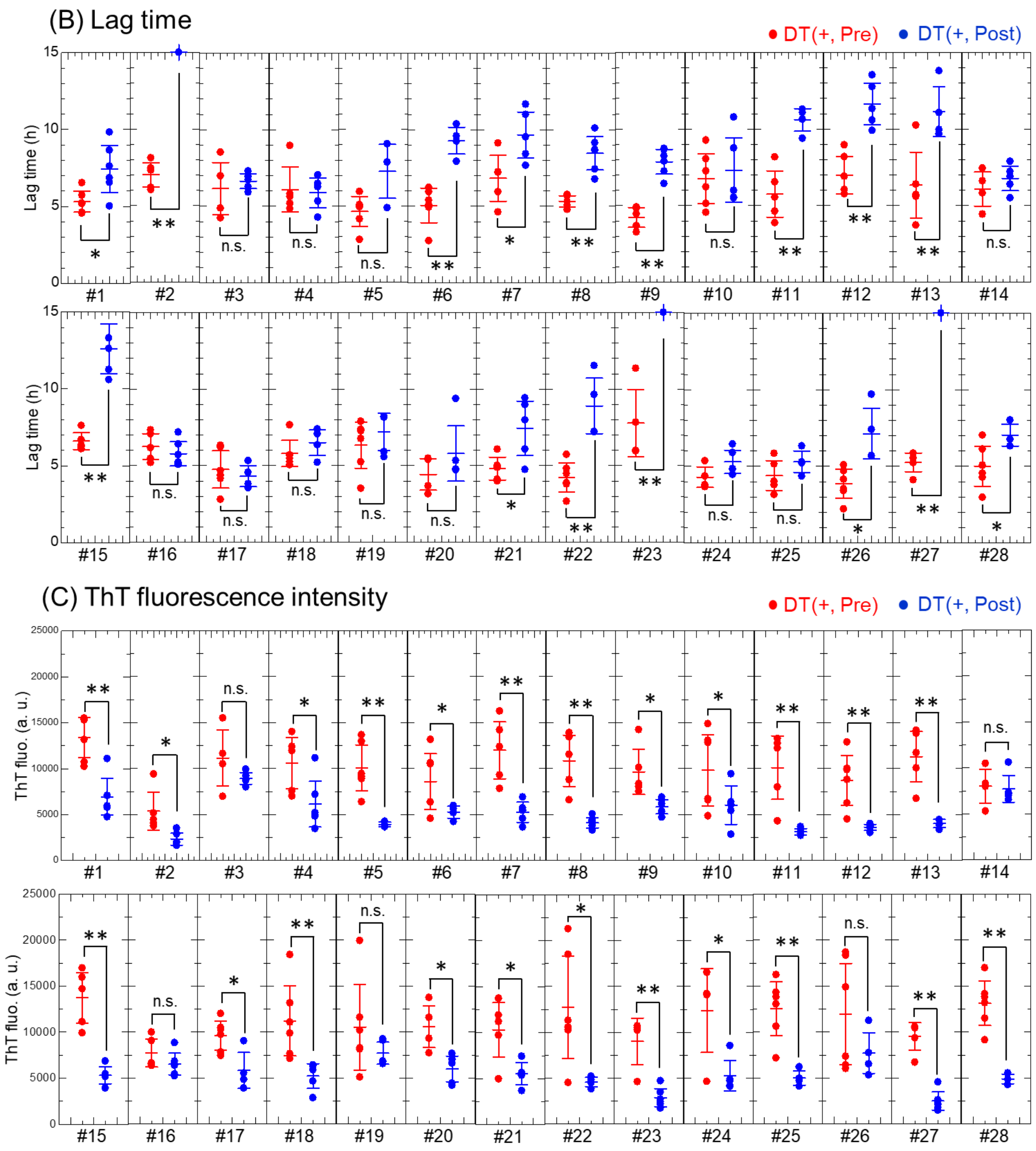


**Figure S8.** (A) ThT kinetics of amyloid fibril formation using sera from dialysis patients ($N$ = 28) collected immediately before (red curves) and after (blue curves) maintenance dialysis treatments. Test of significance of (B) lag time and (C) ThT fluorescence intensity of the HANABI assay using sera collected before and after maintenance dialysis treatment. The symbols **, *, and n.s. correspond to the $p$-values of $p < 0.01$, $0.01 < p < 0.05$, and $p > 0.05$, respectively. The $p$-value was calculated by the unpaired $t$-test.

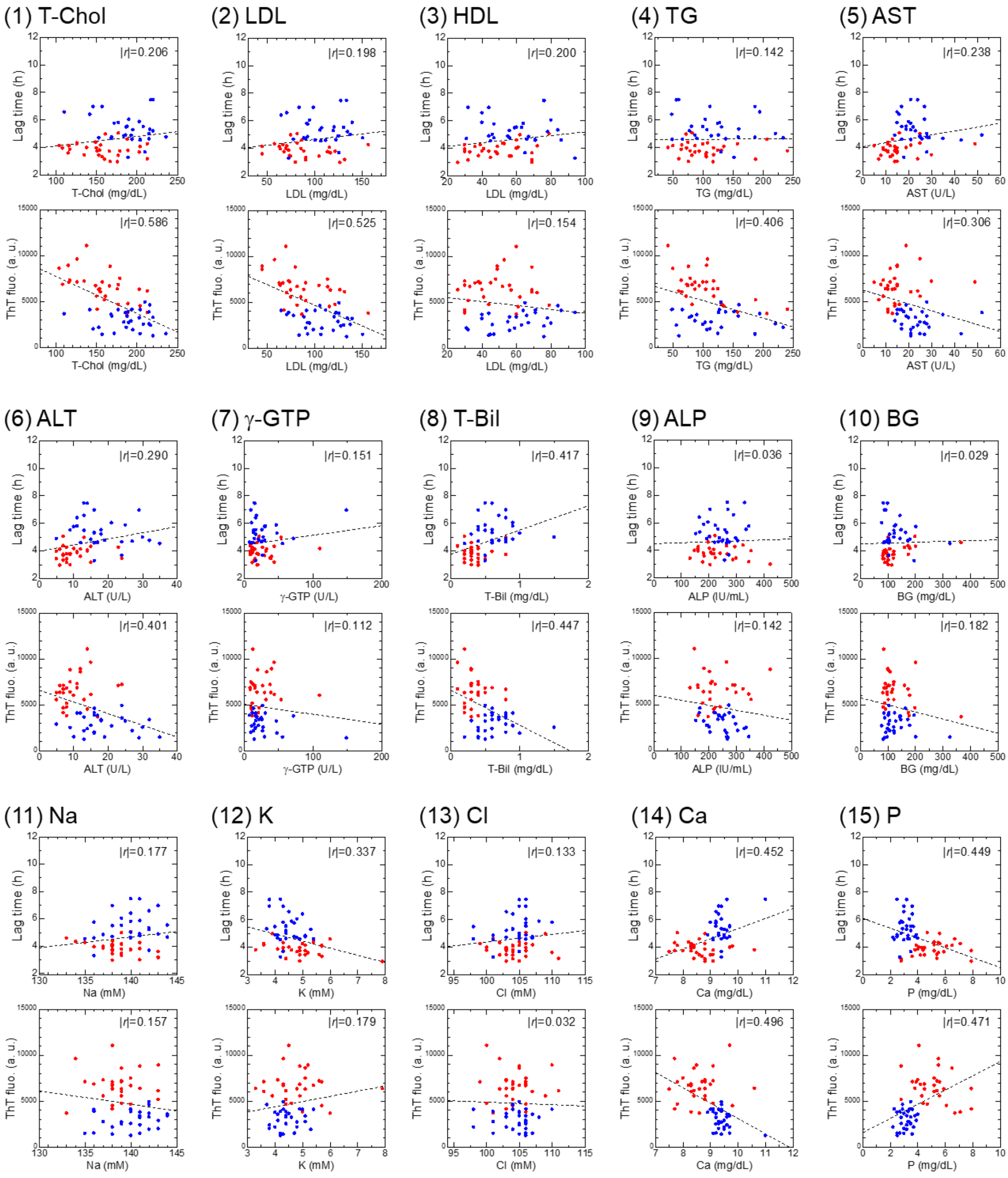


(Continue to the next page)

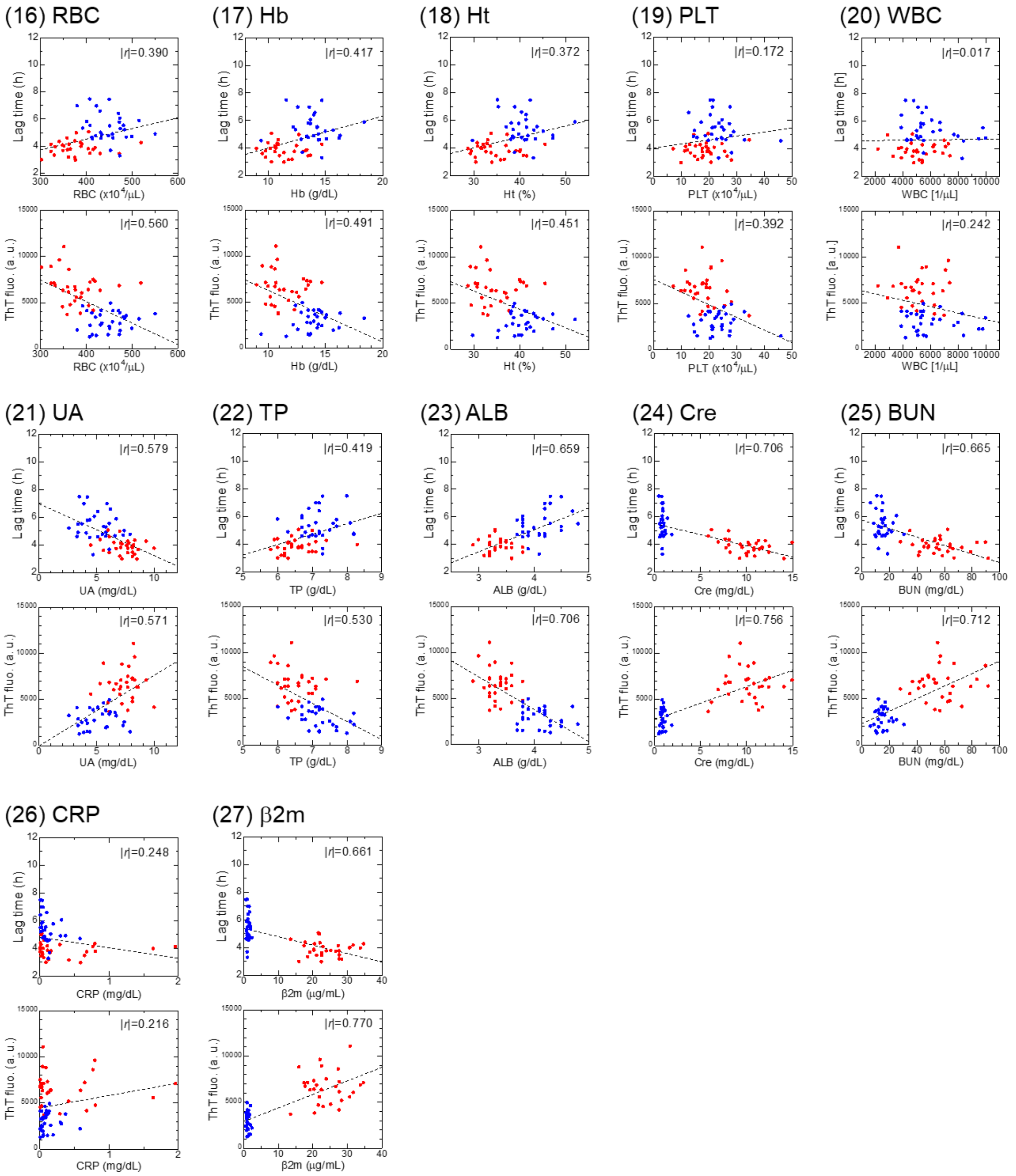


**Figure S9.** Analysis of correlation coefficients between 27 serum components and results of ThT fluorescence kinetic analysis. For each component, data were fitted by a linear function. Abbreviations of serum components are as follows: (1) T-Chol, total cholesterol; (2) LDL, low-density lipoprotein; (3) HDL, high-density lipoprotein; (4) TG, triglyceride; (5) AST, aspartate acid transaminase; (6) ALT, alanine aminotransferase;

(7) γ-GTP, γ-glutamyl transpeptidase; (8) T-Bil, total bilirubin; (9) ALP, alkaline phosphatase; (10) BG, blood glucose; (16) RBC, red blood cell; (17) Hb, hemoglobin; (18) Ht, hematocrit; (19) PLT, platelet; (20) WBC, white blood cell; (21) UA, ureic acid; (22) TP, total protein; (23) ALB, serum albumin; (24) Cre, creatinine; (25) BUN, blood urea nitrogen; and (26) CRP, C-reactive protein.

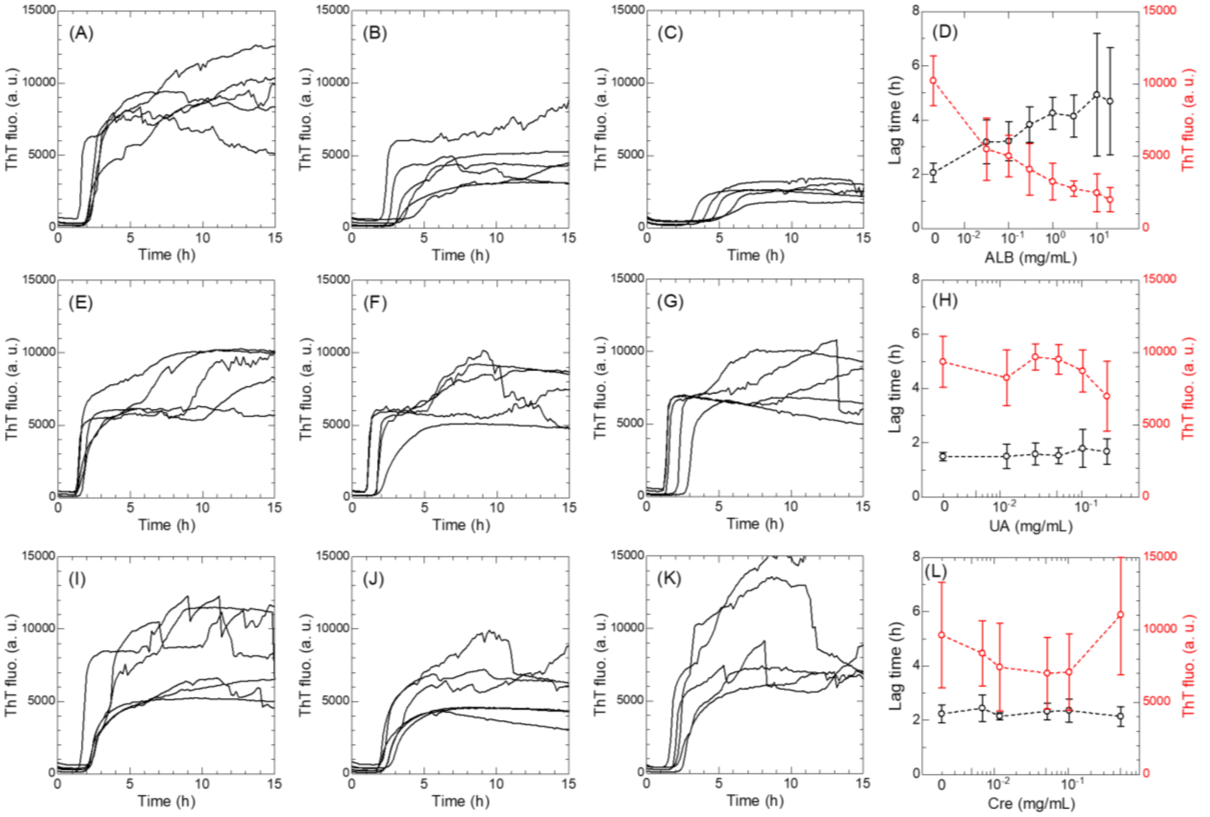


**Figure S10.** Effects of (A-D) serum albumin (ALB), (E-H) ureic acid (UA), and (I-L) creatinine (Cre) on amyloid fibril formation of recombinant β2m monomers. ThT kinetics of samples including serum albumin with concentrations of (A) 0, (B) 0.03, and (C) 3 mg/mL; ureic acid with concentrations of (E) 0, (F) 0.01, and (G) 0.1 mg/mL; and creatinine with concentrations of (I) 0, (J) 0.05, and (K) 0.5 mg/mL. Dependency of the lag time and ThT fluorescence on (D) serum albumin, (E) ureic acid, and (L) creatinine concentrations are summarized. Error bars denote the standard deviation. For each concentration, measurements were performed using independent sample solutions (*n* = 5).

**Table S1.** Summary of the examination of significance shown in Figure S7B,C.

| | Lag time | ThT fluo. |
|---|---|---|
| $p > 0.05$ (n.s.) | 12 | 5 |
| $0.01 < p < 0.05$ (*) | 5 | 10 |
| $p < 0.01$ (**) | 11 | 13 |

**Table S2.** Summary of correlation-coefficient analysis for identical serum components for the lag time.

| Rank | Serum component | Correlation-coefficient |
|---|---|---|
| 1 | Cre | 0.706 |
| 2 | BUN | 0.665 |
| 3 | β2m | 0.661 |
| 4 | ALB | 0.659 |
| 5 | UA | 0.579 |
| 6 | Ca | 0.452 |
| 7 | P | 0.449 |
| 8 | TP | 0.419 |
| 9 | Hb | 0.417 |
| 10 | T-Bil | 0.417 |

**Table S3.** Summary of correlation-coefficient analysis for identical serum components for the ThT fluorescence

| Rank | Serum component | Correlation-coefficient |
|---|---|---|
| 1 | β2m | 0.770 |
| 2 | Cre | 0.756 |
| 3 | BUN | 0.712 |
| 4 | ALB | 0.706 |
| 5 | T-Chol | 0.586 |
| 6 | UA | 0.571 |
| 7 | RBC | 0.560 |
| 8 | TP | 0.530 |
| 9 | LDL | 0.525 |
| 10 | Ca | 0.496 |

#### Supplementary References


1 Hoshino, J. *et al.* Carpal tunnel surgery as proxy for dialysis-related amyloidosis: results from the Japanese society for dialysis therapy. *Am J Nephrol* **39**, 449-458, doi:10.1159/000362567 (2014).

2 Hoshino, J. *et al.* Significance of the decreased risk of dialysis-related amyloidosis now proven by results from Japanese nationwide surveys in 1998 and 2010. *Nephrol Dial Transplant* **31**, 595-602, doi:10.1093/ndt/gfv276 (2016).

3 Jarrett, J. T. & Lansbury, P. T., Jr. Seeding "one-dimensional crystallization" of amyloid: a pathogenic mechanism in Alzheimer's disease and scrapie? *Cell* **73**, 1055-1058, doi:0092-8674(93)90635-4 (1993).

4 Shahnawaz, M. *et al.* Development of a Biochemical Diagnosis of Parkinson Disease by Detection of alpha-Synuclein Misfolded Aggregates in Cerebrospinal Fluid. *JAMA Neurol* **74**, 163-172, doi:10.1001/jamaneurol.2016.4547 (2017).

5 Das, A. & Mukhopadhyay, C. Urea-mediated protein denaturation: a consensus view. *J Phys Chem B* **113**, 12816-12824, doi:10.1021/jp906350s (2009).

6 Zhang, C. M. *et al.* Possible mechanisms of polyphosphate-induced amyloid fibril formation of beta2-microglobulin. *Proc Natl Acad Sci U S A* **116**, 12833-12838, doi:10.1073/pnas.1819813116 (2019).

7 Noji, M. *et al.* Heating during agitation of beta2-microglobulin reveals that supersaturation breakdown is required for amyloid fibril formation at neutral pH. *J Biol Chem* **294**, 15826-15835, doi:10.1074/jbc.RA119.009971 (2019).

8 Noji, M. *et al.* Breakdown of supersaturation barrier links protein folding to amyloid formation. *Commun Biol* **4**, 120, doi:10.1038/s42003-020-01641-6 (2021).

9 Kardos, J., Yamamoto, K., Hasegawa, K., Naiki, H. & Goto, Y. Direct measurement of the thermodynamic parameters of amyloid formation by isothermal titration calorimetry. *J Biol Chem* **279**, 55308-55314, doi:10.1074/jbc.M409677200 (2004).

10 Auer, S. & Frenkel, D. Prediction of absolute crystal-nucleation rate in hard-sphere colloids. *Nature* **409**, 1020-1023, doi:10.1038/35059035 (2001).

11 Guo, Z., Jones, A. G. & Li, N. The effect of ultrasound on the homogeneous nucleation of BaSO4 during reactive crystallization. *Chemical Engineering Science* **61**, 1617-1626, doi:10.1016/j.ces.2005.09.009 (2006).

12 Chiba, T. *et al.* Amyloid fibril formation in the context of full-length protein: effects of proline mutations on the amyloid fibril formation of beta2-microglobulin. *J Biol Chem* **278**, 47016-47024, doi:10.1074/jbc.M304473200 (2003).

13 Nakajima, K. *et al.* Optimized sonoreactor for accelerative amyloid-fibril assays through enhancement of primary nucleation and fragmentation. *Ultrason Sonochem* **73**, 105508, doi:10.1016/j.ultsonch.2021.105508 (2021).

14 Nakajima, K. *et al.* Nucleus factory on cavitation bubble for amyloid beta fibril. *Sci Rep* **6**, 22015, doi:10.1038/srep22015 (2016).

15 Hirota-Nakaoka, N., Hasegawa, K., Naiki, H. & Goto, Y. Dissolution of beta2-microglobulin amyloid fibrils by dimethylsulfoxide. *J Biochem* **134**, 159-164, doi:10.1093/jb/mvg124 (2003).

16 Ogi, H. Wireless-electrodeless quartz-crystal-microbalance biosensors for studying interactions among biomolecules: a review. *Proc Jpn Acad Ser B Phys Biol Sci* **89**, 401-417, doi:10.2183/pjab.89.401 (2013).

17 Noi, K., Iwata, A., Kato, F. & Ogi, H. Ultrahigh-Frequency, Wireless MEMS QCM Biosensor for Direct, Label-Free Detection of Biomarkers in a Large Amount of Contaminants. *Anal Chem* **91**, 9398-9402, doi:10.1021/acs.analchem.9b01414 (2019).

18 Katou, H. *et al.* The role of disulfide bond in the amyloidogenic state of beta(2)-microglobulin studied by heteronuclear NMR. *Protein Sci* **11**, 2218-2229, doi:10.1110/ps.0213202 (2002).

19 Lee, W., Tonelli, M. & Markley, J. L. NMRFAM-SPARKY: enhanced software for biomolecular NMR spectroscopy. *Bioinformatics* **31**, 1325-1327, doi:10.1093/bioinformatics/btu830 (2015).

20 Ikenoue, T. *et al.* Heat of supersaturation-limited amyloid burst directly monitored by isothermal titration calorimetry. *Proc Natl Acad Sci USA* **111**, 6654-6659, doi:1322602111/10.1073/pnas.1322602111 (2014).

21 Nakajima, K. *et al.* Half-Time Heat Map Reveals Ultrasonic Effects on Morphology and Kinetics of Amyloidogenic Aggregation Reaction. *ACS Chem Neurosci* **12**, 3456-3466, doi:10.1021/acschemneuro.1c00461 (2021).

22 Gejyo, F., Homma, N., Suzuki, Y. & Arakawa, M. Serum levels of beta 2-microglobulin as a new form of amyloid protein in patients undergoing long-term hemodialysis. *N Engl J Med* **314**, 585-586, doi:10.1056/NEJM198602273140920 (1986).

23. Gejyo, F., (2019) Dialysis-related amyloidosis -from the beginning of the research to the future- In Retire. Fumitake Gejyo from Pres. Niigata Univ.
